# Antisense transcription-dependent chromatin signature modulates sense transcription and transcript dynamics

**DOI:** 10.1101/187237

**Authors:** Thomas Brown, Françoise S. Howe, Struan C. Murray, Emily Seward, Scott Rata, Andrew Angel, Jane Mellor

## Abstract

Antisense transcription is widespread in genomes. Despite large differences in gene size and architecture, we find that yeast and human genes share a unique, antisense transcription-associated chromatin signature. We asked whether this signature is related to a biological function for antisense transcription. Using quantitative RNA-FISH, we observed changes in sense transcript distributions in nuclei and cytoplasm as antisense transcript levels were altered. To determine the mechanistic differences underlying these distributions, we developed a mathematical framework describing transcription from initiation to transcript degradation. At *GAL1*, high levels of antisense transcription alters sense transcription dynamics, reducing rates of transcript production and processing, while increasing transcript stability, which is also a genome-wide association. Establishing the antisense transcription-associated chromatin signature through disruption of the Set3C histone deacetylase activity is sufficient to similarly change these rates even in the absence of antisense transcription. Thus, antisense transcription alters sense transcription dynamics in a chromatin-dependent manner.

Graphical Abstract

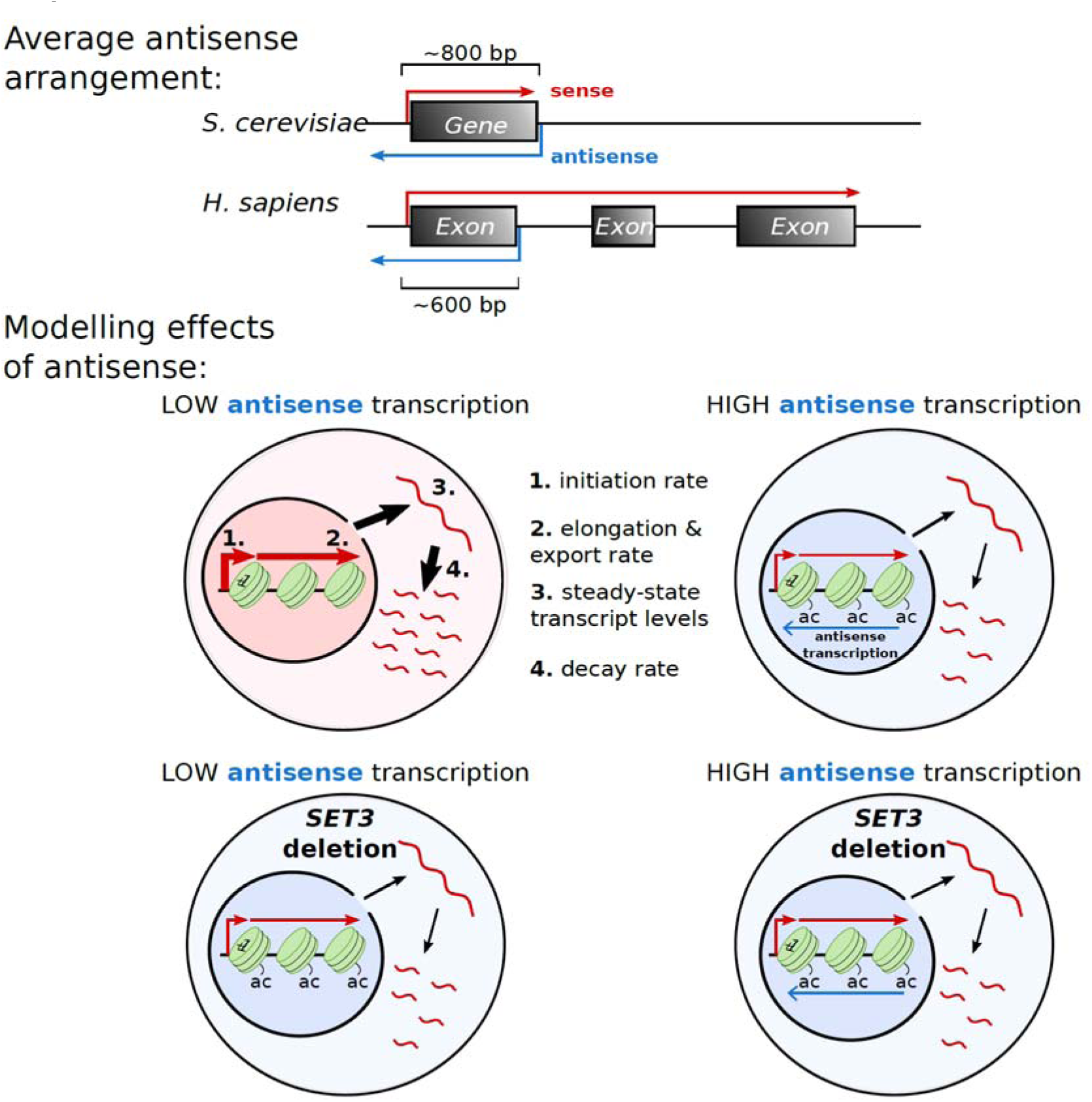

In this work, Brown et al. provide a mechanistic understanding of the effect of antisense transcription on the production and fate of sense transcripts. Antisense transcription buffers genes against the action of the Set3 lysine deacetylase, thus altering rates of transcript production, processing and stability.

- Conserved antisense transcription-dependent chromatin architecture near promoters
- Antisense transcription alters sense transcription dynamics and transcript stability
- Antisense transcription functions in a chromatin-dependent manner
- Increased acetylation by *set3Δ* mimics high antisense transcriptional dynamics

## Introduction

The transcription of genomes is not limited to the transcription of genes alone. Transcription is a universally pervasive and interleaved process, with transcription events initiating from regulatory sequences such as enhancers, divergently from gene promoters, and on the antisense strand of genes (Lam et al., 2014; Mellor et al., 2016; Murray and Mellor, 2016; Tisseur et al., 2011). Nascent transcripts at enhancers or around promoters can be used to recruit (Battaglia et al., 2017) and activate (Bose et al., 2017) epigenetic modifiers associated with chromatin and to activate neighbouring genes in a cell type-specific manner (Werner et al., 2017). This may explain why some chromatin modifications only appear after transcription has initiated (Howe et al., 2017). In addition to transcripts, co-transcriptional processes also influence chromatin modifications in genomes. Transcription of many non-coding transcripts uses a form of RNA polymerase II (RNAPII) that is depleted for conserved features normally associated with efficient transcription elongation including serine 2 phosphorylation of the C-terminal domain (CTD) on the largest RNAPII subunit, the H3K36 methyltransferase Set2 and the elongation factor Paf1 (Fischl et al., 2017; Murray et al., 2015). One such class of non-coding transcripts contains the nascent transcripts transcribed from the antisense strand of the gene, which are often rapidly degraded by exonucleases (He et al., 2008; Neil et al., 2009; van Dijk et al., 2011). Although antisense transcription within genes is a consistent feature of eukaryotic genomes (Mellor et al., 2016), it is not known whether it is simply a by-product of gene transcription, whether there are consequences of antisense transcription and, if so, whether these are conserved across species. Much effort has been expended to determine the function(s) associated with antisense transcription. For a small number of genes, sense and antisense transcription appear to suppress one another and/or be reciprocally regulated (Camblong et al., 2007; Castelnuovo et al., 2013; Hongay et al., 2006; Houseley et al., 2008). However, there is no obvious global relationship between sense and antisense transcription, as levels at the same gene are not correlated genome-wide, either positively or negatively (Murray et al., 2015) and a recent study found that, at the protein level, gene expression is unaffected by lowering levels of antisense transcription in the majority of the 162 genes studied (Huber et al., 2016).

As antisense transcription often proceeds into the sense promoter of its associated gene (Mayer et al., 2015; Murray et al., 2015; Xu et al., 2011) and does not appear to be contemporaneous with sense transcription (Castelnuovo et al., 2013; Nguyen et al., 2014), we previously hypothesised that antisense-transcribing RNAPII might indirectly influence sense transcription by modulating the chromatin environment in the vicinity of the sense promoter. Thus one round of antisense transcription would be sufficient to leave an epigenetic signature and influence sense transcription. We identified in yeast a chromatin signature at the sense promoter and in the early coding region unique to genes with high levels of antisense transcription: high levels of nucleosome occupancy leading to a reduced nucleosome depleted region (NDR), high histone H3 lysine acetylation and histone turnover, but low levels of histone H3 lysine 36 tri-methylation (H3K36me3), H3K79me3 and H2BK123 mono-ubiquitination, amongst others (Murray et al., 2015). Some of these features have been found associated with antisense transcription in mammals (Lavender et al., 2016). Conservation of chromatin features associated with antisense transcription between yeast and mammals will enable us to apply anything we learn in yeast about the mechanistic consequences of antisense transcription more broadly. Here we address the question of how antisense transcription influences sense transcription using a stochastic model of transcription and quantitative data from a single-molecule approach, RNA fluorescence in situ hybridization (RNA-FISH). This allows for the best understanding of the effects of antisense transcription on the dynamics of sense transcript production and processing at the individual cell level. We model RNA-FISH data obtained from engineered constructs expressing high or low levels of antisense transcription, but the same level of sense transcripts, thus mimicking the commonly reported situation where antisense transcription has little effect on steady-state transcript levels. We show that antisense transcription decreases rates of transcript production and processing while increasing transcript stability and, importantly, that these changes in transcription dynamics are directly influenced by the antisense-dependent chromatin signature. As we reveal a remarkably conserved chromatin architecture around the sense promoter and early transcribed region of yeast and human genes with antisense transcription, despite large differences in gene size, we suggest that the effect of antisense transcription is likely to be conserved between yeast and human genes.

## Results

### A conserved arrangement of sense and antisense transcription start sites in yeast and human genes

To address whether and how antisense transcription is conserved across species, it was necessary to map genic transcription start sites (TSSs) as either *sense* sites (sTSS), or *antisense* sites (asTSS), depending on their orientation relative to their proximal gene, and the extent of sense and antisense transcription downstream of these TSSs (Figure 1A). As many antisense transcripts are unstable, we used data from nascent transcript mapping techniques such as NET-seq (Churchman and Weissman, 2011; Nojima et al., 2015), PRO-seq (Booth et al., 2016) or GRO-seq (Core and Lis, 2008) to assess genome-wide levels of transcription in *S.cerevisiae* and HeLa cells. To map TSSs, we used Cap Analysis of Gene Expression (CAGE) data for HeLa cells (Consortium et al., 2014), pooling the polyadenylated and non-polyadenylated tag data from nuclear, cytoplasmic and whole cell fractions, and TIF-seq for yeast (Pelechano et al., 2014). It is important to note that we may fail to identify the TSSs of unstable antisense transcripts and thus underestimate actual levels of antisense transcription. From over 20,000 protein-coding genes we identified 9,320 with a sTSS in HeLa cells. Of these genes, we found 2,468 (27%) with an internal asTSS. 1,008 (40%) of these asTSSs were within 500 bp of the sTSS, with a median distance of 632 bp (Figure 1B). Thus, a large fraction of active genes in HeLa cells show evidence of a productive, antisense-oriented transcription start site close to their promoter. We defined 5,222 yeast genes with a sTSS, of which 1,529 (29%) had an asTSS. The median distance between the sTSSs and asTSSs of yeast genes was 884 bp, 252 bp larger than in humans (Figure 1C).

**Fig 1.**
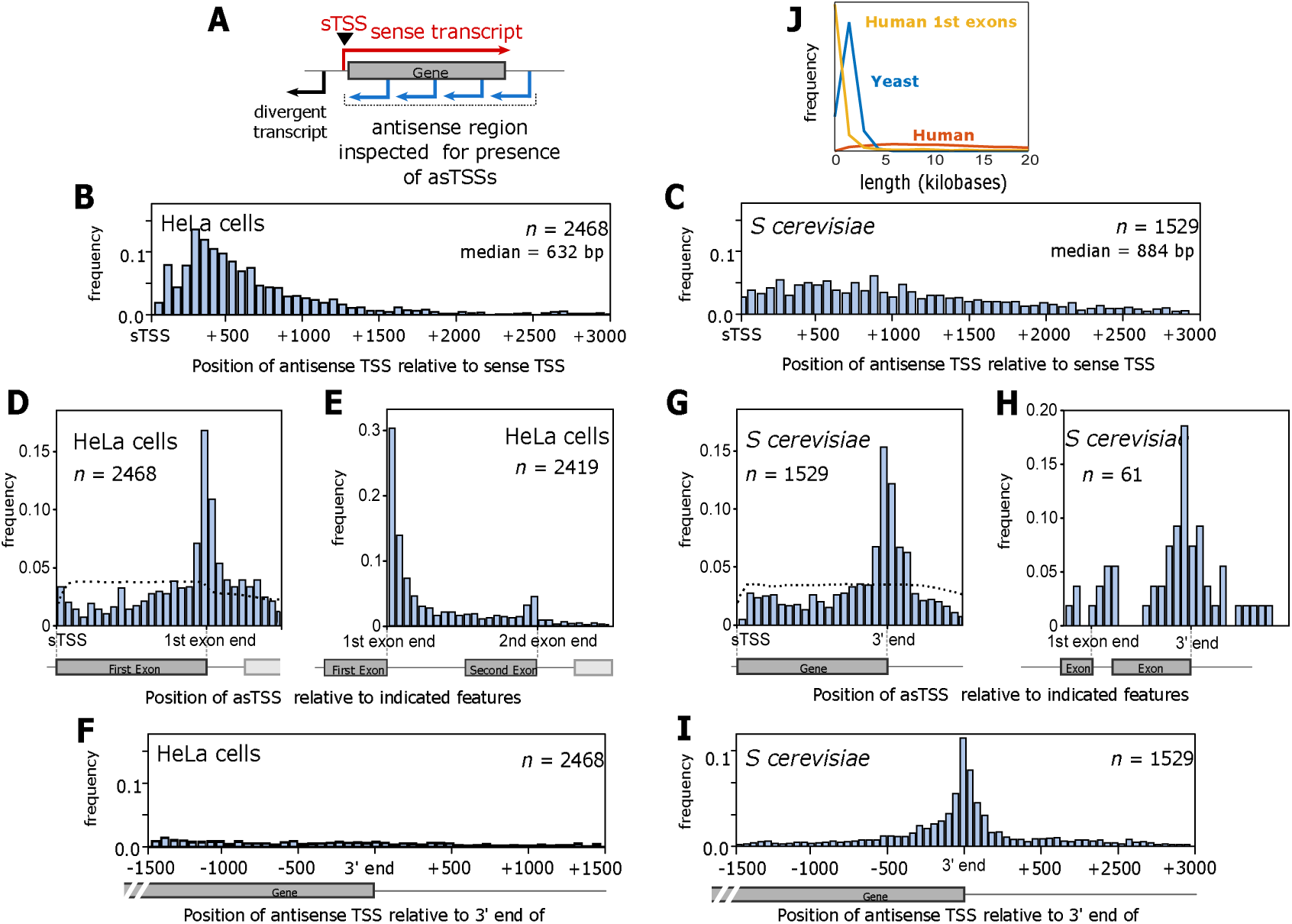
Antisense transcripts initiate at a similar distance from the sense TSS in yeast and humans, though from a distinct functional site. **(A)** A schematic demonstrating how antisense TSSs (asTSSs) were defined in this study. Blue arrows represent possible sites of antisense transcript initiation, within the region inspected for the presence of asTSSs, as defined by CAGE. **(B)** The distribution of distances between sTSS and asTSS in HeLa cells, for those 2,468 genes which had both an upstream sTSS and an internal asTSS defined by CAGE. Shown is the median distance between the sTSS and asTSS. **(C)** The distribution of distances between the sTSS and asTSS in *S. cerevisiae* (budding yeast), for those 1,529 genes that both an overlapping sense and antisense transcript, defined by TIF-seq. **(D)** The position of the asTSS relative to the sTSS *and* the end of the 1^st^ exon in HeLa cells, to demonstrate which of the two points the asTSS aligns to preferentially. This position was defined as the distance between the sTSS and the asTSS, divided by the distance between the sTSS and the end of the 1^st^ exon. The genes are the same as those shown in **B**. The dotted line represents the average distribution from a thousand simulations, in which asTSSs for each gene were randomly reassigned to a base pair within the region shown. **(E)** The position of the asTSS relative to the sTSS and the end of the 2^nd^ exon, for those HeLa genes in **D** that also had a second exon. **(F)** The distribution of distances between the 3’ end of genes and asTSS in HeLa cells, for the same genes in **B. (G)** The position of the asTSS relative to the sTSS and the 3’ end of the open reading frame of the 1,529 *S. cerevisiae* genes that have both an overlapping sense and antisense transcript. The dotted line was generated as in **D. (H)** The position of the asTSS relative to the end of the 1^st^ exon and the 3’ end of the open reading frame of those *S. cerevisiae* genes in **G** that also have an intron. **(I)** The distribution of distances between the 3’ end of *S. cerevisiae* genes and asTSS in, for the same genes in **C. (J)** The distribution of lengths for human 1^st^ exons, human genes, and yeast genes.

Strikingly, in humans, the asTSS aligned more closely to the 1^st^ exon-intron boundary than the sTSS, and with a much higher frequency than expected if asTSSs are randomly redistributed over this region (Figure 1D). In fact, in 2,162 (88%) genes with antisense transcripts, the asTSS is closer to the 1^st^ exon-intron boundary than it is to the sTSS. Using the same approach, we also found that the asTSS aligned much more tightly with the 1^st^ exon-intron boundary than it did with the 2^nd^ exon-intron boundary (Figure 1E) or to the 3’ end (Figure 1F). In yeast, antisense transcription tends to initiate from the vicinity of the 3’ region of genes (Figure 1G) (Xu et al., 2011) rather than from introns (Figure 1H). 1,202 (79%) genes with an antisense transcript had an asTSS closer to their 3’ end than to their sTSS (Figure 1I). Despite their distinct sites of origin in humans and yeast (1^st^ intron-exon boundary and 3’ end respectively; Figures 1D,G,I) and gene size (Figure 1J), the asTSS is at a similar median distance to the sTSS in humans and yeast (632bp compared to 884 bp), suggesting a conserved arrangement.

### Higher nucleosome occupancy at promoters of genes with high antisense transcription in yeast and humans

To examine how antisense transcription influences the chromatin and sense transcription in the vicinity of the promoter, we assessed three regions: 300 nucleotides upstream of the sTSS (the sense promoter), 300 nucleotides downstream of the asTSS (the antisense promoter), and the region between the two TSSs, which was broken into an equal number of bins. We compared the upper and lower quintiles of genes with an asTSS, giving us two groups of 494 genes in humans, and 306 and 307 genes in budding yeast. Firstly we compared levels and distributions of sense and antisense transcription in the three regions using NET-seq, GRO-seq or PRO-seq, which are similar when comparing HeLa cells with yeast and the different techniques (Figure 2A). From this point, in Figures 2 and 3, data are related to profiles for NET-seq, while the PRO-seq and GRO-seq profiles are in supplemental figures 1 (relates to Figure 2) and 2 (relates to Figure 3). Next, we examined a genome-wide map of nucleosome occupancy (MNase-seq). Both species had a nucleosome-depleted region at the asTSS and a marked *increase* in nucleosome occupancy in the vicinity of the sTSS in genes with the highest levels of antisense transcription (Figure 2B). This suggests that antisense transcription may modulate promoter chromatin in both species *without* necessarily altering levels of sense transcription in the vicinity of the sense promoter (Figure 2C,D). Indeed, nucleosome occupancy and sense transcription may well be disconnected (Nocetti and Whitehouse, 2016), despite the apparent association between sense transcription and nucleosome depletion at the promoter (Figure 2B). Nucleosomes also interact with one another in local space, one component of the chromatin conformation in the nucleus (Hsieh et al., 2015). To see if the antisense-associated increase in nucleosome occupancy is related to the chromatin conformation, we used Micro-C data in yeast, which identifies chromosomal contacts at the resolution of nucleosomes (Hsieh et al., 2015). For the 5,222 yeast genes that had an associated sense transcript as described above, we determined its level of gene *compaction*, as defined by Hsieh et al., (2015), normalised such that it is independent of gene length. Strikingly, we found an inverse association between intragenic contacts and antisense transcription. Genes with an antisense transcript showed a significantly reduced level of gene compaction (p = 1.2×10^−65^, Wilcoxon rank sum test, Figure 2E), suggesting that antisense transcription favours a looser higher order structure. For sense transcription there was *no* significant change in the level of compaction (p = 0.29, Wilcoxon rank sum test, Figure 2E).

**Fig 2.**
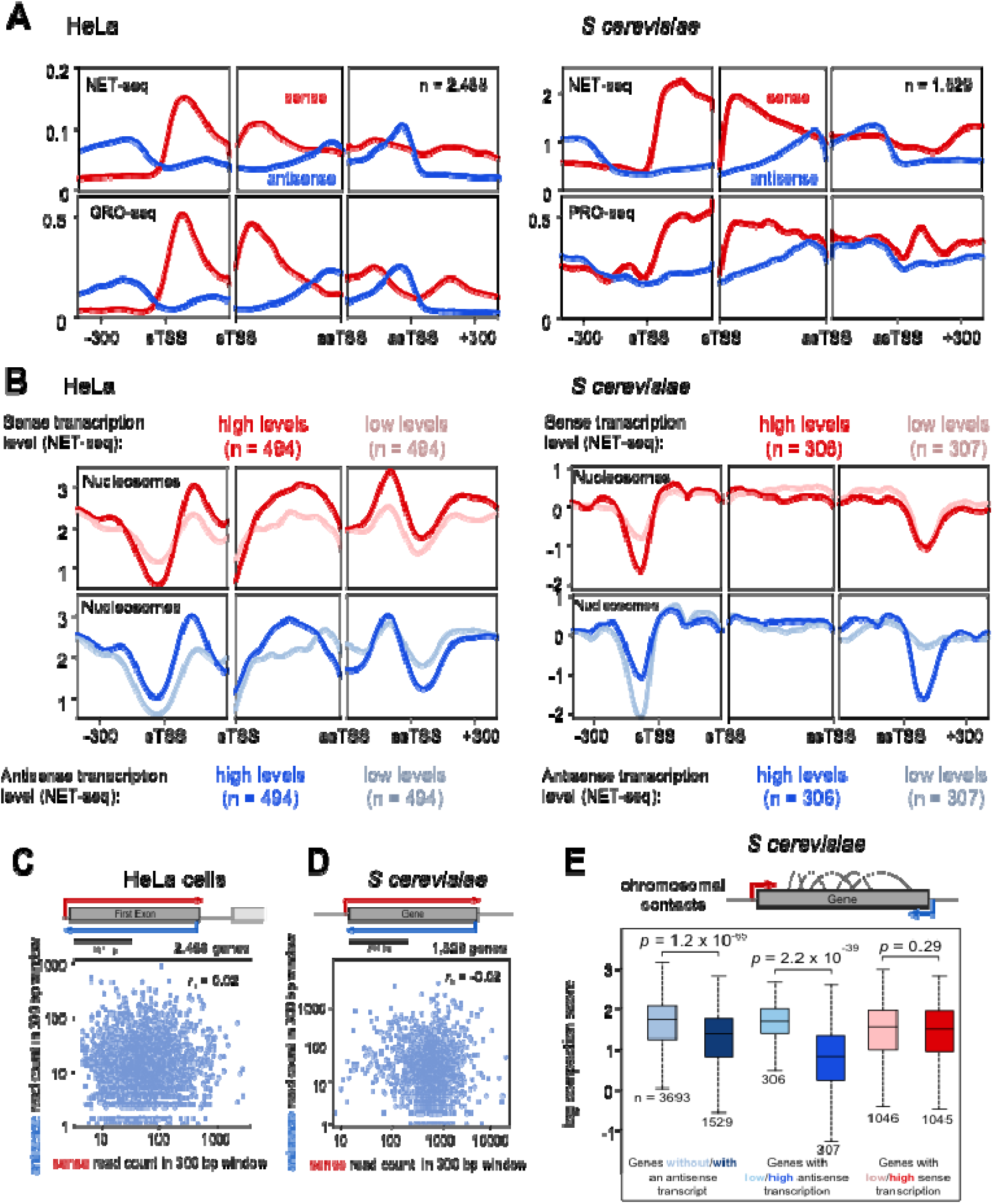
Antisense transcription is associated with changes in chromatin structure in both yeast and humans, but not with changes to the level of sense transcription. **(A)** The average levels of sense and antisense transcription determined for both HeLa and *S. cerevisiae* using NET-seq, HeLa using GRO-seq, and *S. cerevisiae* using PRO-seq. For each trio of panels, the left panel shows average levels around the sTSS, the right panel shows the average levels around the asTSS, and the middle panel shows the average level within thirty equal sized bins within the region bound by the sTSS and asTSS. In all cases, levels of transcription on the sense strand are shown in red, while levels on the antisense strand are shown in blue. Genes considered are those which contained an asTSS, as defined in Fig 1. **(B)** The average levels of nucleosome occupancy determined for both HeLa and *S. cerevisiae* using MNase-seq. Panels are grouped in threes and show average levels as in **A**. The top panels compare two sets of genes – those with high levels of sense transcription (dark red), and those with low levels (pale red). The bottom panels show those genes with high levels of *antisense* transcription (dark blue), and those with low levels (pale blue). **(C-D)** Scatter plots comparing the number of sense and antisense NET-seq reads within the 300 bp window shown, in both HeLa cells and *S. cerevisiae*, for those genes with both an sTSS and asTSS. Shown for both species is the Spearman’s correlation coefficient, *r*_*s*_. **(E)** Boxplots showing the distribution of gene compaction in different sets of *S cerevisiae* genes. Gene compaction was determined by summing the number of intragenic contacts, measured by Micro-C, and dividing by gene length. The numbers at the bottom of each box plot show the number of genes in that group.

**Fig 3.**
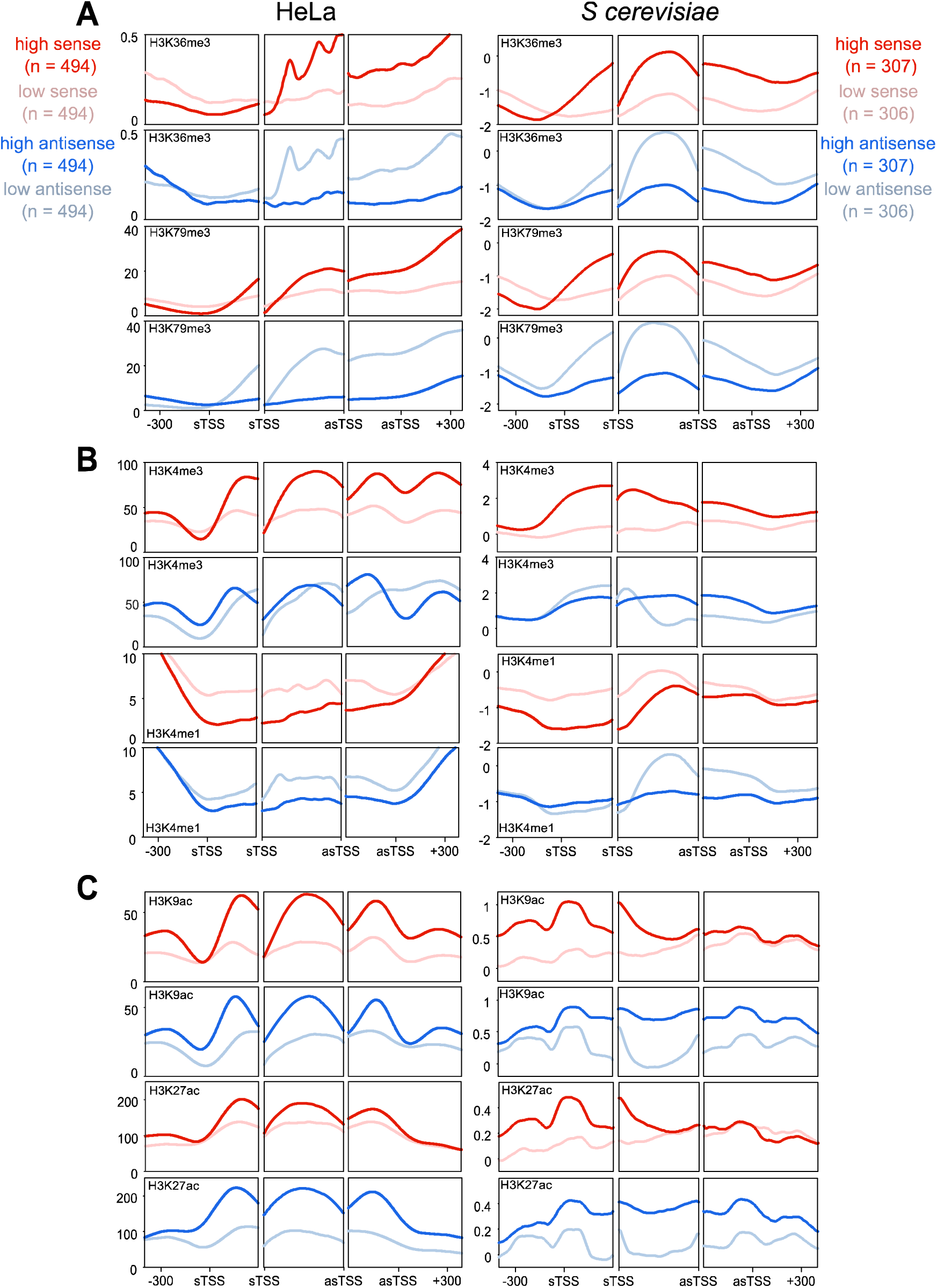
Antisense transcription has similar associations with chromatin modifications in both yeast and humans. **(A)** The average levels of H3K36me3 and H3K79me3 and antisense transcription in HeLa and *S. cerevisiae* genes. For each trio of panels, the left panel shows average levels around the sTSS, the right panel shows the average levels around the asTSS, and the middle panel shows the average level within thirty equal sized bins within the region bound by the sTSS and asTSS. Genes considered are selected from those which contained an asTSS, as defined in Fig 1. Shown in red are two sets of genes – those with high levels of sense transcription (dark red), and those with low levels (pale red). Shown in blue are those genes with high levels of *antisense* transcription (dark blue), and those with low levels (pale blue). **(B)** Average levels of H3K4me3 and H3K4me1, laid out as in **A. (C)** Average levels of H3K9ac and H3K27ac, laid out as in **A**.

### Antisense transcription is associated with a similar unique chromatin signature in yeast and humans

We next turned our attention towards histone modifications (Figure 3, NET-seq and Supplemental Figure 2, PRO-seq and GRO-seq). Levels of H3K36me3 and H3K79me3 were higher in the region bounded by the sTSS and asTSS for those genes with high *sense* transcription, in both species (Figure 3A) (Ng et al., 2003; Pokholok et al., 2005). Strikingly, however, these two modifications are much lower in those genes with high levels of *antisense* transcription, in both humans and yeast (Figure 3B). This is despite the fact that the level of sense transcription is the same in the high/low antisense classes – i.e. there is no difference in gene *activity* (see Figure 2C,D). That these modifications should have reverse associations with sense and antisense in *both* yeast and humans is intriguing, and suggests there may be some fundamental difference to the two modes of transcription that is shared across species. By contrast, levels of H3K4me3 tended to be more evenly spread between the sTSS and the asTSS in genes with high antisense transcription compared to high sense transcription (Figure 3B). Levels of H3K4 lysine mono-methylation (H3K4me1) tended to be lower with high sense or antisense transcription showing that not all modifications have reciprocal patterns with sense or antisense transcription. Finally, levels of H3 acetylation are increased in the presence of antisense transcription (Figure 3C), particularly in the region downstream of the sense TSSs.

Taken all together, one can see that despite the vast differences in size between yeast and human genes, they share a very similar arrangement in terms of where their antisense transcripts initiate relative to their coding-transcript start site, and in how antisense transcription associates with numerous shared chromatin features. We conclude that antisense transcription in the vicinity of the sense promoter is associated with increased histone lysine acetylation and nucleosome occupancy, and decreased histone H3K36me3, H3K79me3 and chromatin compaction, and that this unique architecture is conserved between yeast and humans.

Is there a consequence of this widespread and conserved antisense transcription initiating downstream from the sense promoter for the genes in yeast and human that have it? How might it be changing gene behaviour? To address this, we developed a mathematical model that describes the dynamics of transcription and allows us to discriminate between transcriptional events at the sense promoter, the nucleus and the cytoplasm. When compared with experimental data, the model allows us to determine which parameters of sense transcription production and processing are affected by antisense transcription.

### A stochastic model for transcription

Our stochastic model of transcription captures the production, processing and destruction of a transcript (Figure 4A) and builds on existing models of transcription (Choubey et al., 2015; Raj et al., 2006; Zenklusen et al., 2008). Within the model, a gene promoter is allowed to switch between an active and inactive state stochastically, with an activation rate α and an inactivation rate β. In the active state, transcription initiation occurs with rate γ. As a result, the “Mean Production rate”, i.e. the average rate of transcript initiation, is given by 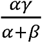

**Fig 4.**
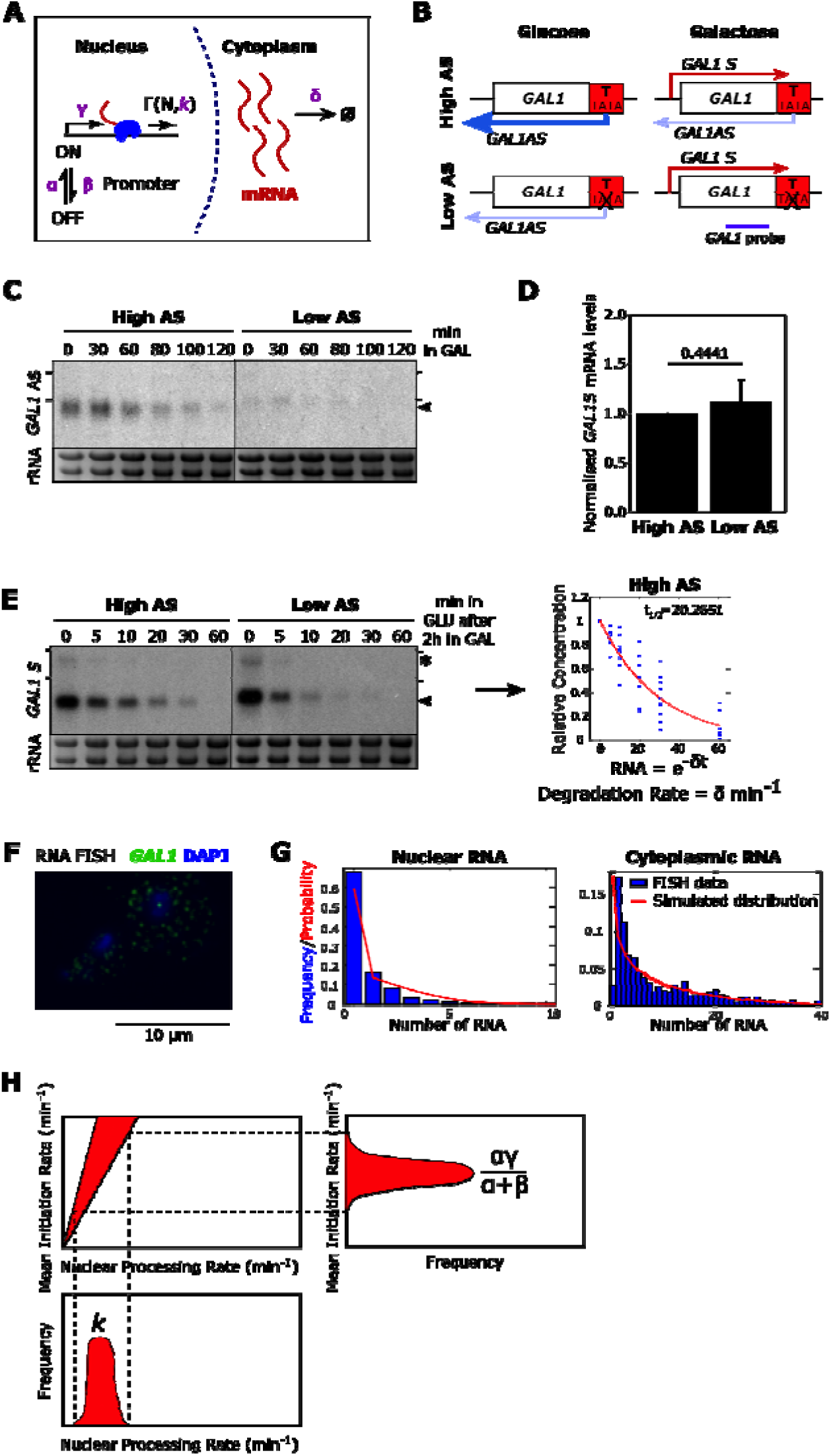
A stochastic model for transcription dynamics. **(A)** Schematic of the model (see text for details). **(B)** The engineered *GAL1* expressing high or low antisense (AS) transcription (Murray et al., 2015). Purple line shows the position of the strand-specific probes used for Northern blotting. Sense transcripts are in red, antisense transcripts in blue. **(C)** Representative Northern blot showing levels of *GAL1* antisense transcripts (black arrowhead) in the high and low antisense strains during the transition from glucose (0 min) to galactose (GAL). RNA-FISH experiments are performed after 120 min in GAL. Samples were run on the same gel with the intervening lanes spliced out (as indicated by the black vertical line). The positions of the 25S and 18S rRNAs are represented by short black horizontal lines. Ethidium bromide-stained rRNA is the loading control. **(D)** Quantitation of *GAL1* sense transcripts levels as measured by Northern blotting in the high and low antisense strains, normalised to High AS levels. N=9, error bars are SD, *p-*value shown above the bars. **(E)** Representative Northern blot showing *GAL1* sense transcripts (black arrowhead) and *GAL10* lncRNA (asterisk) after transfer from GAL (0) to GLU for the time indicated (min). From these data the rates of degradation of the transcripts are calculated, after normalization to the GLU timepoint. Samples were run on the same gel with the intervening lanes spliced out (as indicated by the black vertical line). The positions of the 25S and 18S rRNAs are represented by short black horizontal lines. N= 9. **(F)** Example of single molecule RNA-FISH data showing two cells. DNA is stained with DAPI (blue) and single *GAL1* sense transcripts in green. A bright nuclear focus is present in the top cell, containing 2-3 nascent transcripts. **(G)** The frequency of nuclear and cytoplasmic transcripts for 1193 individual cells averaged across 9 experiments are determined using an automated foci recognition algorithm (blue bars). The red line shows the simulated distribution from the model in **(A)**. Data shown are counts and fit for Low AS *GAL1* mRNA. **(H)** Schematic showing how mean initiation rate and nuclear processing rates are obtained. The fit to nuclear RNA distribution dictates the ratio of mean initiation rate to nuclear processing rate and the cytoplasmic rate determines the ratio of mean initiation rate to degradation rate. By fitting the degradation rate to the shutdown data **(E)**, the probability distributions for mean initiation rate and nuclear processing rate are obtained outright.

Nuclear transcript processing, meanwhile, is modelled as a sum of reactions representing the advancement of RNAPII across the DNA as a series of stochastic jumps. The time to fully process a nuclear transcript is distributed as a sum of *N* exponentials, corresponding to gene length, with parameter *k* corresponding to elongation rate, Γ(*N,k*). Our experimental protocol does not allow for the separation of nascent and nuclear transcripts, therefore in our modelling framework we do not differentiate between the two types of transcripts. As a result, the parameter *k* is a conflation of both elongation rate and nuclear export rate. We refer to this parameter as the “Nuclear Processing rate”, representing the time for a transcript to go from initiation to export. Finally, transcripts are assumed to degrade in the cytoplasm with a constant half-life, decaying exponentially with Degradation rate δ.

We used this model to study the transcription dynamics of the inducible yeast gene *GAL1* (Figure 4B). The first strain (High AS) contains an engineered form of the *GAL1* gene that expresses a stable antisense transcript as a result of insertion of the *ADH1* transcription terminator *(GAL1∷ADH1t)* (Murray et al., 2015; Murray et al., 2012). In the second strain (Low AS), a 6 bp AT rich sequence within the inserted terminator region is scrambled (while retaining the overall base composition), resulting in a significant reduction in levels of the antisense transcript (Figure 4C), which is a consequence of reduced levels of antisense transcription (Murray et al., 2015), but no change in levels of sense transcripts (Figure 4D).

Two types of data were used to parameterize the dynamics of transcription. Firstly, we obtained the rate of degradation of cytoplasmic sense transcripts (Figure 4E), relying on the galactose-inducible and glucose-repressible nature of *GAL1*. We grew cells in galactose-containing media, then recorded the decreasing concentration of sense mRNA via Northern Blot at multiple timepoints after cells were moved to glucose-containing media. By fitting an exponential curve to the datasets, we obtained the Degradation rate δ, which gives the half-life via the formula: 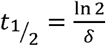

Secondly, we obtained the distribution of individual sense transcripts in the nucleus and cytoplasm of cells using RNA-FISH (Figure 4F). We probed cells grown in galactose-containing media for 2h for the *GAL1* sense transcript and counted the number of fluorescent foci within the nucleus and cytoplasm. Individual dots were assumed to represent at least one transcript, with the number of transcripts at a given dot determined by dividing the intensity of the dot by the median intensity of all foci. Several hundred of cells were considered for a given experiment and distributions of nuclear and cytoplasmic transcript counts were obtained (Figure 4G). The nuclear distribution gives an indication of Mean Initiation rate relative to Nuclear Processing rate, or the fraction 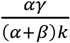. The cytoplasmic distribution, correspondingly, tells us the Mean Initiation rate relative to Degradation rate, or 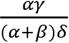

Using the measured Degradation rate **5**, we find the parameters that best fit the RNA-FISH data via the Kolmogorov-Smirnov test, sampling 1,000,000 parameter sets via Latin Hypercube (McKay et al., 1979) and sampling the 1,000 parameter sets with the best Kolmogorov-Smirnov Statistic. The parameters obtained by simulation are then used to determine the Mean Initiation rate 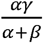 and Nuclear Processing rate *k*. The 1,000 best parameter sets obtained from the cytoplasmic data give a probability distribution for the expected true value of the parameter. The nuclear data show the likely corresponding ratio of Mean Initiation rate to Nuclear Processing rate. By projecting the probability distribution for the Mean Initiation rate onto the linear relationship, we obtain a probability distribution for the Nuclear Processing rate (Figure 4H). Taking the most likely value from the probability distribution, we obtain the inferred parameter values for the given strain. The various steps involved in generating rates for initiation of transcription (min^−1^) and nuclear processing rate (min^−1^) are shown in Figure 4H. Therefore, for any strain we can obtain the Mean Initiation rate, Nuclear Processing rate and Degradation rate, corresponding to promoter, nuclear and cytoplasmic effects on sense transcript dynamics (Supplemental Figure 3).

### Antisense transcription influences rates of transcription initiation, nuclear processing and transcript stability

We generated experimental data using the two strains in which *GAL1* was subject to different levels of antisense transcription (Figure 4B). By obtaining transcriptional parameters in these two strains we were able to compare how antisense transcription influences sense transcription and transcripts. Strikingly, the stability of the engineered *GAL1* sense transcripts is higher with greater antisense transcription (t_1/2_ = 13.53 vs. 20.26 min for low vs high AS; Figures 4E, 5B). Furthermore, the effect of antisense transcription on transcript stability is not strictly limited to the engineered *GAL1* genes used here. Comparing the stability of transcripts produced from 1,529 yeast genes with, and 3,693 without, an antisense transcript revealed a significant (*p* = 8 x 10^−13^) increase in stability for sense transcripts produced from genes with an antisense transcript, using stability data obtained from Wang et al. 2002 (Wang et al., 2002) (Figure 5A).

**Fig 5.**
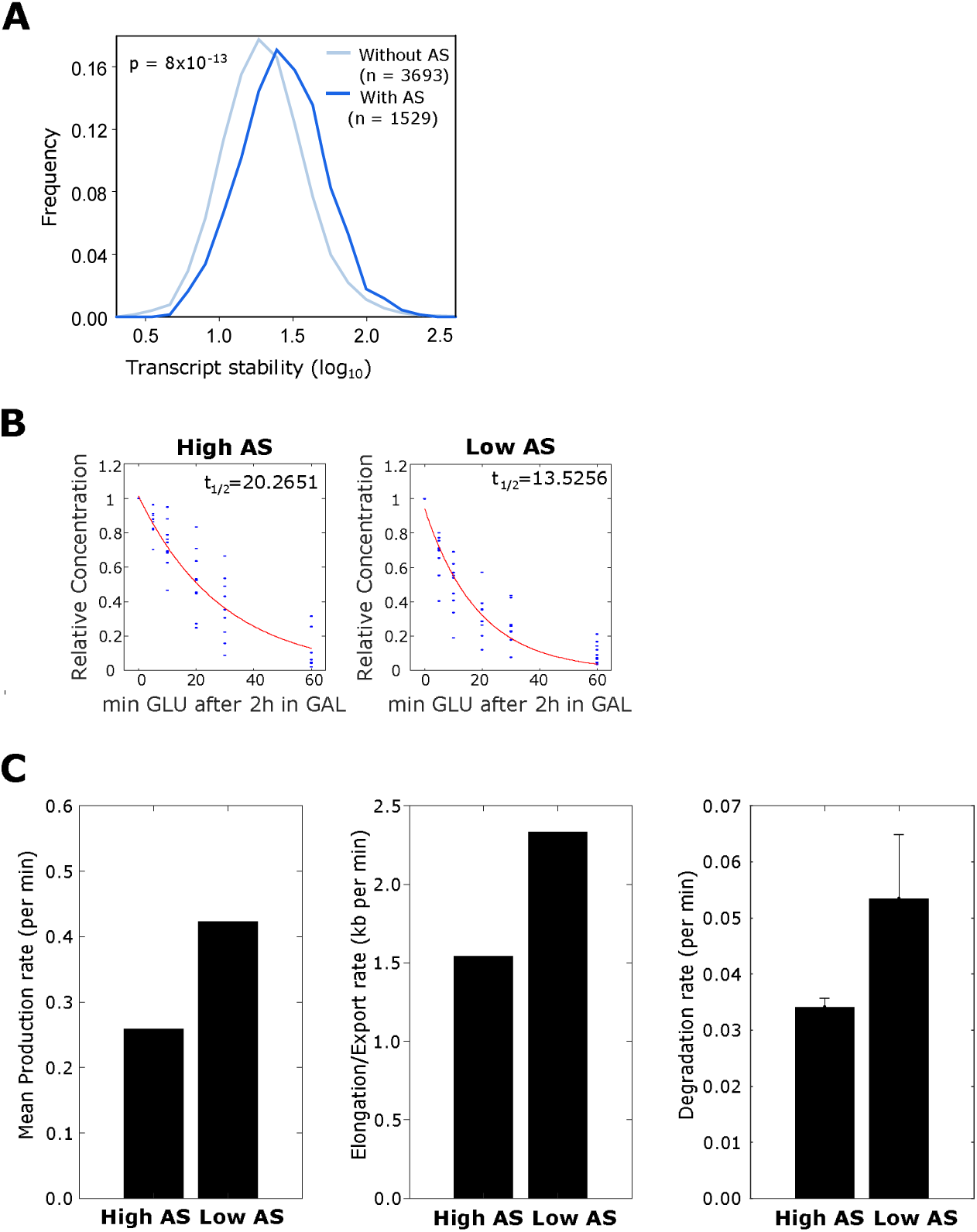
Transcription dynamics change with high antisense transcription. (**A**) Transcript stability for sense transcripts from 1,529 genes with an asTSS (dark blue) or 3,693 genes without one (light blue). The frequency plot shows the majority of transcripts have higher stability when expressed from genes with an antisense transcript. (**B**) *GAL1* sense transcript turnover rates from the engineered *GAL1* with high (left) or low (right) antisense transcription. The red lines show the best fit for 9 experiments (see Figure 4E for details). (**C**) Histograms showing the Mean Production rate (left), Mean Elongation/Export rate (middle) and Transcript Degradation rate (right; error bars for degradation rates are RMSE of linear regression fit to exponential model) for the engineered *GAL1* genes expressing high or low antisense transcription.

Modelling of the RNA-FISH data reveals roles for antisense transcription in controlling the rates of initiation and nuclear processing of *GAL1* transcription/transcripts; both parameters were lower in the construct expressing higher antisense transcription (Figure 5B). The Mean Production rate was 0.425 min^−1^ in the construct with low antisense transcription, but 0.256 min^−1^ in the construct with high antisense transcription. Similarly, the Nuclear Processing (Elongation/Export) rate was reduced from 2.33 min^−1^ to 1.54 min^−1^ in the presence of antisense transcription. Taken together, these data suggest compensating changes in rates of sense transcript production and sense transcript degradation as a result of antisense transcription. In the case of the engineered *GAL1* gene, antisense transcription does not alter overall sense transcript levels but *does* alter the dynamics of sense transcript production. It is possible that changes in transcription dynamics resulting from antisense transcription are a consequence of altered patterns of histone modification. To this end, we sought to assess whether experimentally modulating histone modifications could recapitulate the effects of changing antisense transcription, focusing on histone H3 lysine acetylation. Genes with high levels of antisense transcription tend to be associated with increased histone acetylation compared to genes with lower levels. The histone deacetylase complexes Rpd3L, Rpd3S and Set3C lead to increased global acetylation and control transcript dynamics at a small number of genes, but do not affect global gene expression (Kim et al., 2016; Kim et al., 2012; Pijnappel et al., 2001; Woo et al., 2017). We asked whether the HDACs, by their effect on histone acetylation (increased levels by virtue of ablating the deacetylase activity), might modulate the same parameters as antisense transcription.

### *SET3* deletion differentially influences levels of H3K9ac at genes that differ by the presence or absence of antisense transcription

H3K9 acetylation levels increase globally in strains lacking specific components of the Set3C, Rpd3L or Rpd3S complexes – *SET3* (encoding Set3, present in Set3C), *PHO23* (in Rpd3L), *HOS2* (in Set3C and Rpd3L) and *RPD3* (in Rpd3S and Rpd3L) (Weinberger et al., 2012). By reanalyzing H3K9ac ChIP-seq data in strains lacking these components (Weinberger et al., 2012), we found that loss of either Set3C (*set3*δ) or Rpd3L (*pho23*δ) resulted in the largest average change in levels of H3K9ac over genes (Supplemental Figure 4A). Next, we asked whether the change in H3K9ac, following gene deletion, is different when considering those genes with or without an antisense transcript (Supplemental Figures 4B,C). Strikingly, following deletion of *SET3* (and so the Set3 complex), those 3,693 genes *without* an antisense transcript show a significantly larger increase (*p* = 6 x 10^−13^) in the level of H3K9ac than those 1,529 genes *with* an antisense (Figure 6A). No such difference was observed following deletion of *PHO23* (and so the Rpd3L complex) between genes with and without an antisense (Supplemental Figure 4C). We hypothesized that the reason for this large difference following *SET3* deletion is that, with respect to H3K9ac, there is some redundancy in the effect of deleting *SET3* and in the effect of antisense transcription.

**Fig 6.**
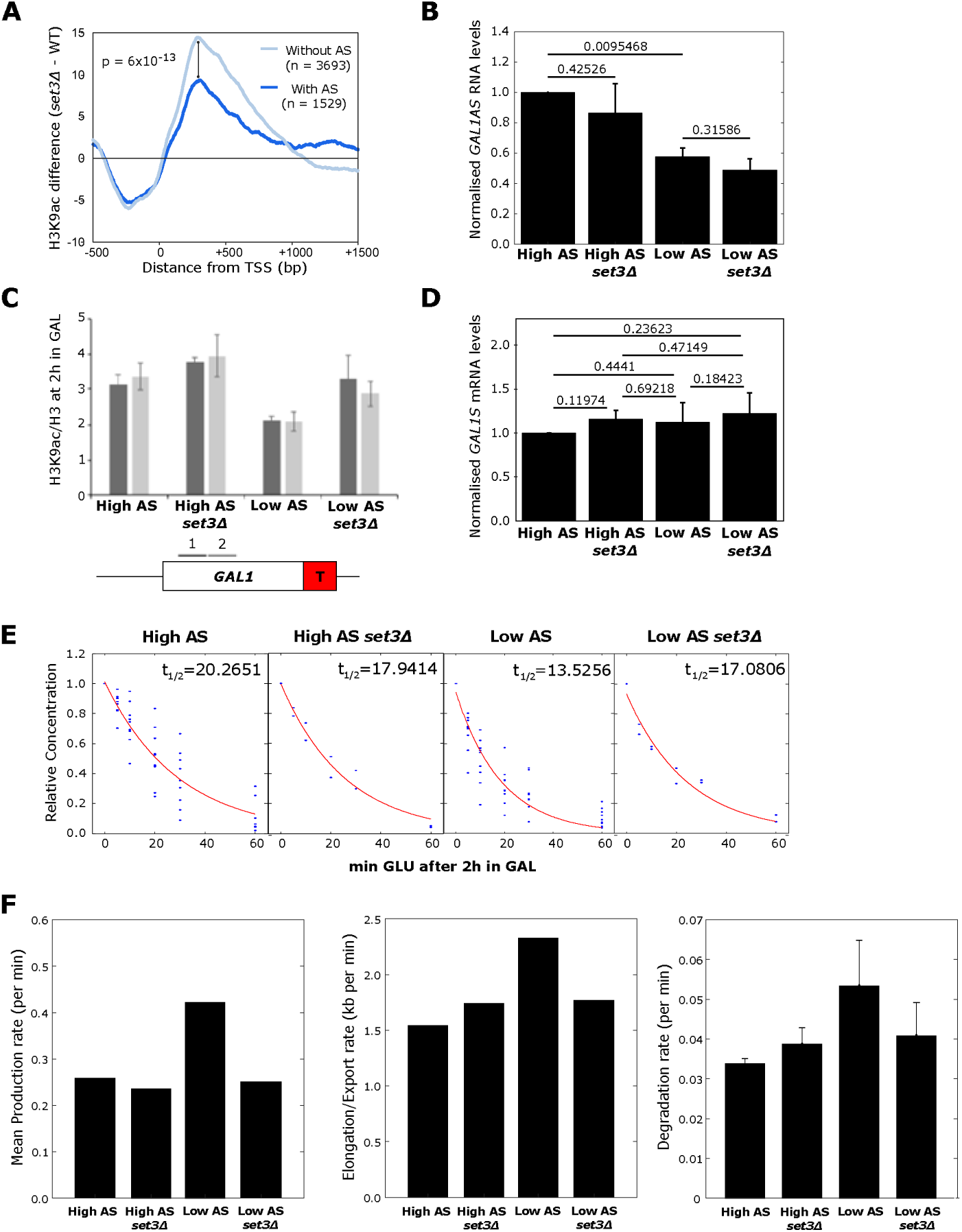
Set3C alters transcription dynamics in an antisense-dependent manner. **(A)** Strains lacking *SET3* show a larger increase in H3K9ac, relative to WT levels, at 3,693 genes without antisense transcripts compared to 1,529 genes with antisense transcripts. **(B)** Quantitation of *GAL1* antisense transcript levels from the high and low antisense constructs with or without *SET3*. N = 2, error bars are SD, *p*-values are shown above the bars. **(C)** Chromatin immunoprecipitation showing levels of H3K9ac relative to histone H3 in the strains with high or low antisense transcription in the presence or absence of *SET3*. The positions of the primer pairs for RT-qPCR are shown in the schematic below. N = 2, error bars are SEM. **(D)** Quantitation of *GAL1* sense transcripts from Northern blotting of the high and low antisense constructs with or without *SET3*. N = 3 error bars are SD. **(E)** *GAL1* sense transcript degradation rates in *SET3* (N = 9) and *set3*Δ (N = 2) strains with high or low *GAL1* antisense transcripts. **(F)** Histograms showing the Mean Production rate (left), Mean Elongation/Export rate (middle) and Transcript Degradation rate (right) for the engineered *GAL1* genes expressing high or low antisense transcription in the presence or absence of *SET3*. Error bars for degradation rates are RMSE of linear regression fit to exponential model.

To explore this potential redundancy, and to assess the extent to which an increase in acetylation is responsible for antisense-dependent changes in transcription dynamics, we investigated the effect of *SET3* deletion in the presence or absence of antisense transcription at the engineered *GAL1*, again using RNA-FISH data to parameterize sense transcription dynamics (Supplemental Figure 3). Importantly, Set3C does not affect the levels of the stable antisense transcript at engineered *GAL1* (Figure 6B). That there is no change in antisense transcription is confirmed by levels of transcription-associated histone modifications H3K4me2 and H3K4me3 in the four strains, which are low without antisense transcription and higher with antisense transcription, and, importantly, do not change when *SET3* is deleted (Supplemental Figure 5).

Next, we asked how Set3C influences levels of H3K9ac at engineered *GAL1* with high or low antisense transcription (Figure 6C). High levels of acetylated H3K9 in chromatin may *result* from, but is not causally related to, histone turnover (Ferrari and Strubin, 2015) and antisense transcription is associated with high levels of histone turnover and H3K9ac (Murray et al., 2015). As expected, the construct with high antisense transcription has higher levels of H3K9ac than the strain with low levels of antisense transcription, despite both constructs producing similar levels of the *GAL1* sense transcript when induced (Murray et al., 2015) (Figure 6D). On deletion of *SET3* we observe a higher level of H3K9ac in the transcribed region in strains with low antisense transcription (Figure 6C). This neatly reproduces what we have observed genome-wide – that *SET3* deletion has a greater effect on acetylation levels in the absence of antisense transcription.

We conclude that the effect of *SET3* deletion on levels of H3K9 acetylation at the engineered *GAL1* gene is unlikely to result from changes to antisense transcription, but from a direct effect on the chromatin. We hypothesise that antisense transcription buffers chromatin against the modulating effects of Set3C during sense transcription. Thus, following deletion of *SET3*, we would expect the transcription dynamics in the strain with low levels of antisense transcription to resemble those in the strain with high antisense transcription, assuming transcription dynamics are influenced solely by the chromatin.

### Altering levels of histone acetylation recapitulates the effect of antisense transcription on sense transcription dynamics

RNA-FISH data (Supplemental Figure 3) and degradation rates (Figure 6E) were produced for *SET3* deletion strains expressing the engineered *GAL1* gene with either high or low levels of antisense transcription. The experimental data were then modelled to estimate the parameters of sense transcription dynamics (Figure 6F). Consistent with our hypothesis, deletion of *SET3* in the strain with low levels of antisense transcription decreased the Mean Production rate, decreased the Nuclear Processing (Elongation/Export) rate and increased the stability of the mature *GAL1* sense transcripts, making the transcription dynamics of the strain with low levels of antisense transcription behave more like the strain with high antisense transcription. Global levels of transcripts are buffered by opposing rate changes for synthesis and degradation resulting in no overall change, as observed previously (Dori-Bachash et al., 2011). We suggest that antisense transcription reduces the sensitivity of genes to deacetylation by Set3C, and this influences transcription dynamics. Thus, at *GAL1*, antisense transcription buffers gene expression against the action of chromatin modifiers such as Set3C. We conclude that sense transcription dynamics are variable and can be modulated by histone modifiers, and therefore histone modifications, in the vicinity of the promoter and early part of the coding region.

In summary, we show that antisense transcription has a conserved spatial and chromatin architecture in both yeast and human genes, focused around the sense promoter and early transcribed region. Modelling with quantitative data reveals that antisense transcription at the engineered *GAL1* locus alters all measurable aspects of sense transcription by decreasing the rates of initiation and processing of the nuclear transcripts, and the cytoplasmic degradation rate.

## Discussion

Antisense transcription is a widespread feature of both yeast and humans genomes. In this work we provide insights into the consequences of antisense transcription on chromatin architecture and sense transcription dynamics, showing that antisense transcription reduces rates of sense transcript production, processing and degradation. The antisense transcription-dependent rate changes can be mimicked by strains lacking Set3, a component of the Set3C HDAC complex. The *set3*δ strain with low *GAL1* antisense transcription showed raised levels of H3K9ac at *GAL1* similar to those observed with high antisense transcription and a concomitant lowering of sense transcript production, processing and degradation rates to resemble those in strains with high antisense transcription (Figure 7). This supports the promoter proximal chromatin structure as the functional consequence of antisense transcription.

**Figure 7.**
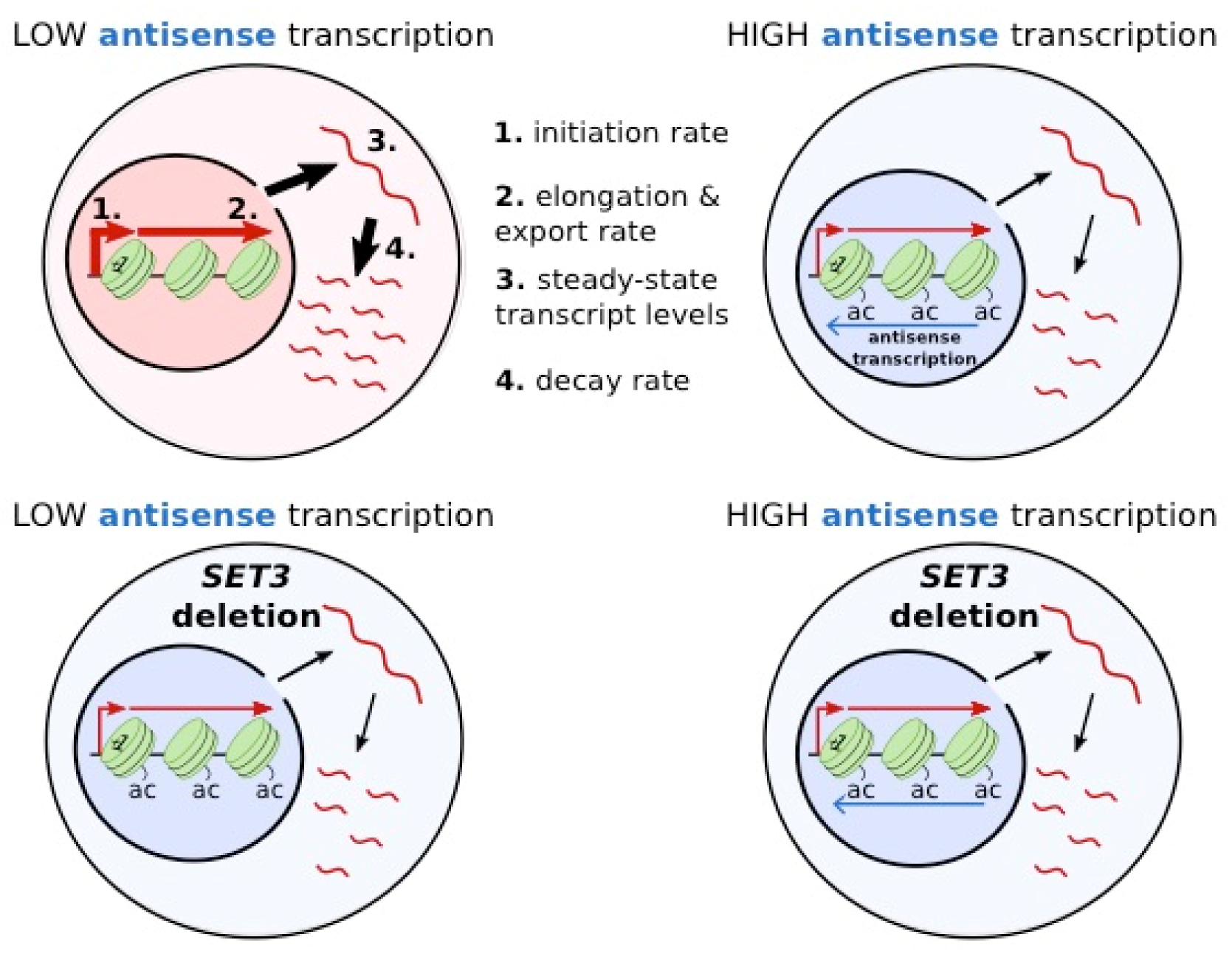
Summary. Schematic outlining the influence of antisense transcription or *SET3* deletion on levels of histone lysine acetylation (ac), steady-state transcript levels (3) and on the rates of transcript production (1), processing and export (2) and transcript decay (4). Line width indicates levels (3) or rates (1,2,4) with increased rate represented by a wider line.

The effects of antisense transcription on the chromatin architecture are conserved between yeast and humans, despite large differences in gene size. In addition, antisense transcription initiates at a similar distance downstream from the sense transcription initiation site in both systems, suggesting that the effect of antisense transcription is focused on the promoter and early-transcribed region of genes. This is where most control can be exerted over transcription dynamics, as the promoter and early coding region chromatin can influence initiation and the elongation phases of transcription respectively. Much transcription initiation in yeast and mammals shows bursting kinetics (Lenstra et al., 2016; Suter et al., 2011). Our model makes no assumptions about the mode of transcription initiation *ab initio* and can accommodate genes with “bursty” kinetics, as it allows for promoters to exist in both active and inactive states, or a constitutive model of production, and so would be applicable to analyse data from mammalian cells. In addition, our model extends previous models, incorporating a stochastic elongation rate, and as such each gene displays a distribution of times from initiation to termination.

At the engineered *GAL1* gene, changing the levels of antisense transcription leads to no detectable difference in steady-state sense transcript levels, although modelling experimental data reveals antisense-dependent reductions in rates of transcript production, processing and degradation. There is a tight, apparently counterproductive, coordination between these processes, with (high) rate of transcription linked to (high) rate of transcript degradation. This coordination is widely observed in yeast and mammals (Das et al., 2017). Here we show for the first time that antisense transcription alters rates of transcription and co-ordinately, rates of transcript degradation. As deletion of *SET3* is sufficient to alter rates from those observed for low antisense at *GAL1* to those for high antisense, with concomitant changes in levels of H3K9ac, we suggest that the effect of antisense transcription is via changes to the promoter proximal chromatin environment. These changes include increased lysine acetylation and increased nucleosome occupancy which together, could influence the residence time of transcription factors bound to the promoter or the composition of RNA polymerase II leaving the promoter. In support of this, we have recently shown that RNA polymerase II shows variable enrichment with elongation factors and that this a function of promoter sequences and associated transcription factors (Fischl et al., 2017). Paf1 enrichment, for example, through its effects on the chromatin structure, affects how the encoded transcripts are decorated with RNA binding proteins that control export from the nucleus. Coordinated transcription and transcript degradation are known to be influenced by promoter sequences, the Rbp4 and 7 components of RNA polymerase II and many factors involved in mRNA degradation (Dori-Bachash et al., 2012; Dori-Bachash et al., 2011; Enssle et al., 1993; Sun et al., 2013) but exactly how antisense transcription via *SET3* (and changes in histone acetylation) interacts with any of these features to change rates remains to be determined. We would favour changes at or around the sense promoter, given the conserved promoter-focused consequence of antisense transcription. In support of this, mutants such as in *RPB4*, which directly reduce rates of transcription, compensate by reducing the rate of transcript degradation (Schulz et al., 2014). Assessment of steady state mRNA levels in such mutants often leads to the conclusion that these factors do not have much effect on gene expression, apart from the associated stress response (O’Duibhir et al., 2014), and this would be entirely consistent with what we observe for antisense transcription and *SET3*.

Could antisense transcription function in gene regulation? By reducing production and increasing stability, as observed with high antisense transcription, the same final transcript response level can be achieved as with low antisense transcription, but the time taken to reach these final levels differs, and this can be a regulatory feature. For example, providing benefit in some bet-hedging strategies (Snijder and Pelkmans, 2011) or if rapidly varying conditions are expected. Indeed, antisense transcription has a proposed role in fine-tuning levels of sense transcripts under different environmental conditions (Xu et al., 2011) and its production can be regulated (Conley and Jordan, 2012; Murray et al., 2015; Murray et al., 2012; Nguyen et al., 2014). Transcription elongation is known to not be a smooth process from start to finish, with RNA polymerase pausing heterogeneously across the gene (Jonkers and Lis, 2015), and this is likely to influence the nuclear processing rate. Changing the nuclear processing rate does not affect the cytoplasmic distribution or the rate at which a cytoplasmic steady-state is reached. However, antisense-transcription-dependent changes to the nuclear processing rate could be tuning how much other pathways can affect a transcript. For example, decreasing the elongation/export rate could make a gene more susceptible to control by a factor that relies on stochastic events during transcription or while a transcript is in the nucleus.

Why does Set3 modulate transcription dynamics predominantly at low antisense genes? Although acetylation levels are generally lower in the low antisense genes, the chromatin for both high and low antisense genes is likely to be dynamically acetylated and deacetylated, as we see increased levels of acetylation when the integrity of Set3C is lost. Interestingly, of the components of HDACs tested, only the strain lacking *SET3* shows a larger increase in acetylation in the chromatin of low antisense genes compared to high antisense genes, both genome-wide and at our engineered *GAL1*. Genes with high antisense show significantly more enrichment for the Spt3 component of the SAGA lysine acetyltransferase (KAT) complex (Murray et al., 2015), supporting a higher inherent level of acetylation at high antisense genes as an explanation for why levels of H3K9ac increase at low antisense genes but not high antisense genes in the *set3Δ* strains. Being able to modulate transcription and transcript dynamics by manipulating the activity of a chromatin modifying enzyme strongly supports antisense transcription modulating the chromatin structure in the vicinity of promoters and this in turn affecting transcription and transcript fate.

Pervasive transcription is not limited to the antisense strand of genes as studied here, but is also abundant at enhancer elements, at the 3’ ends of genes and throughout gene-rich regions of genomes (Andersson et al., 2014; Nojima et al., 2015). In yeast, many non-coding transcription events are mediated with a distinct form of RNA polymerase II, depleted for Paf1 and Set2, amongst other factors (Fischl et al., 2017), explaining in part the unique chromatin environment associated with these events. Whether enhancers also contain a chromatin environment, for example high H3K27ac, dictated by non-coding transcription and how this influences enhancer function are questions for the future, but underscore the potential regulatory nature of non-coding transcription, as distinct from the encoded transcripts. Indeed, it is becoming clear that levels of pervasive transcription can be regulated in genomes (Mellor et al., 2016).

## Author Contributions

Project conception, JM and AA. Experimental work, FSH. Bioinformatics, SCM. Modelling, TB, SR, ES and AA. Image analysis, AA. All authors involved in interpretation of the data. JM and SM wrote the paper with input from FSH, TB and AA.

## Acknowledgements □

We thank the J.M. and A.A. labs for critical discussions, Anitha Nair for excellent technical support, Simon Haenni for 5’ and 3’ RACE mapping of the transcripts, and Ilan Davis and Micron Oxford for microscopy support. This work was supported by: The Wellcome Trust (WT089156MA to J.M.); the BBSRC (BB/P00296X/1 to J.M.); the Leverhulme Trust (RPG-2016-405 to J.M.); a Wellcome Trust Strategic Award (091911) supporting advanced microscopy at Micron Oxford (http://micronoxford.com); EPSRC and BBSRC studentships (EP/F500394/1 to T.B.; EP/G03706X/1 to S.R. BB/J014427/1 to E.S.) and a Royal Society University Research Fellowship (UF120327 to A.A.) J.M. acts as an advisor to and holds stock in Oxford Biodynamics plc., Chronos Therapeutics Ltd., and Sibelius Natural Products Ltd.

---

**Supplemental Figure 1.**
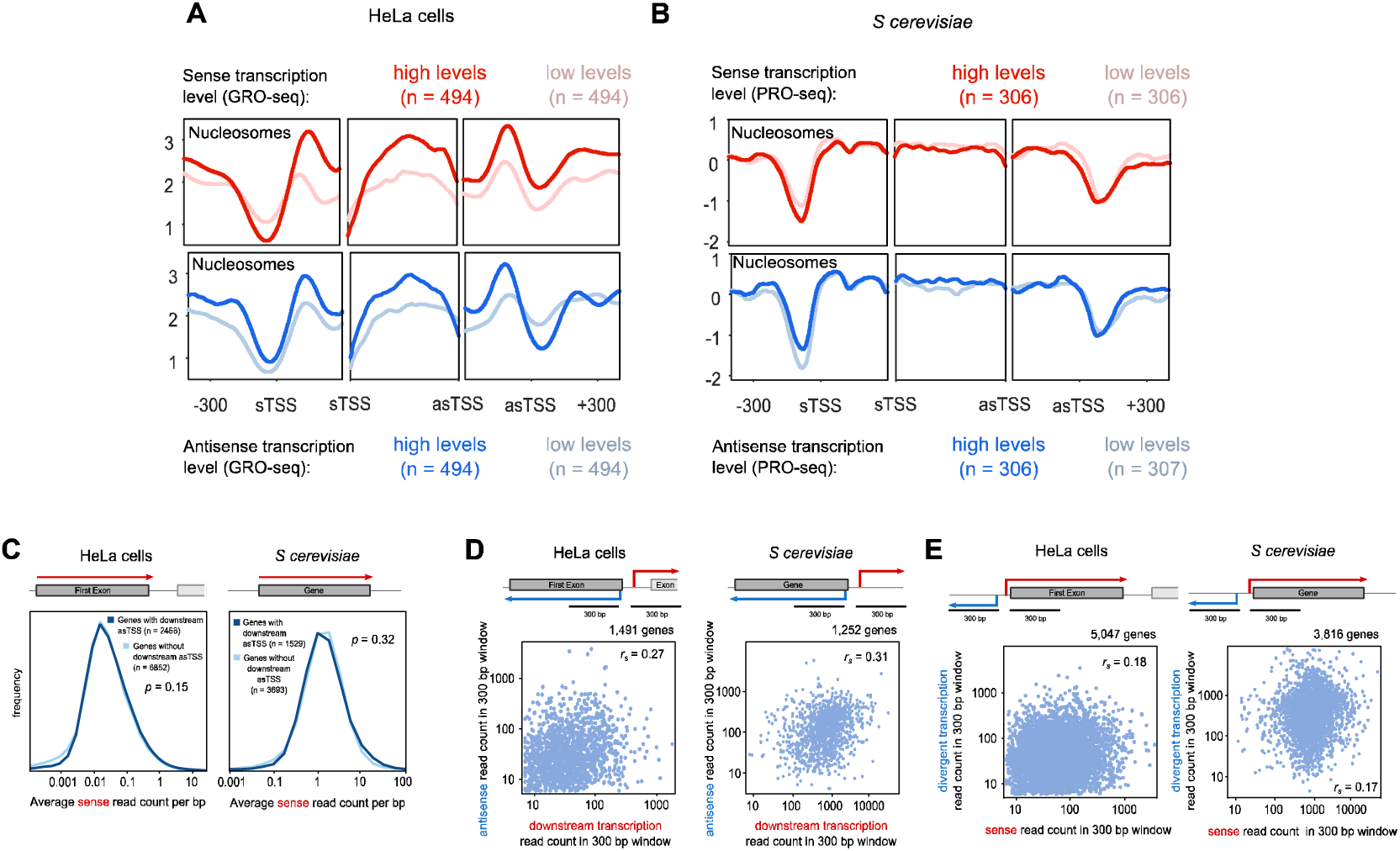
Sense and antisense transcription have similar relationships in yeast and humans. The average levels of nucleosome occupancy determined for both **(A)** HeLa and **(B)** *S. cerevisiae* using MNase-seq. For each trio of panels, the left panel shows average levels around the sTSS, the right panel shows the average levels around the asTSS, and the middle panel shows the average level within thirty equal sized bins within the region bound by the sTSS and asTSS. The top panels compare two sets of genes – those with high levels of sense transcription (dark red), and those with low levels (pale red), determined by GRO-seq in HeLa cells and PRO-seq in yeast. The bottom panels show those genes with high levels of *antisense* transcription (dark blue), and those with low levels (pale blue). **(C)** Distributions of sense transcription reads for genes in HeLa cells and *S. cerevisiae*. Average values per base pair were calculated within the first exon of HeLa cells, and the whole gene of *S. cerevisiae*. Two gene groups were compared – those with an asTSS (dark blue) and those without (light blue). The *p*-value was determined using the Wilcoxon rank sum test. **(D-E)** Scatter plots comparing the number of GRO-seq (HeLa)_or PRO-seq (*S. cerevisiae*) reads at two different windows and orientations, as shown in the gene diagrams. **(D)** Levels of antisense transcription were compared to downstream sense transcription in both HeLa cells and *S. cerevisiae*. **(E)** Levels of sense transcription were compared to upstream divergent transcription in both HeLa cells and *S. cerevisiae*. Shown for both species is the Spearman’s correlation coefficient, *r*_*s*_. Relates to Figure 2.

**Supplemental Figure 2.**
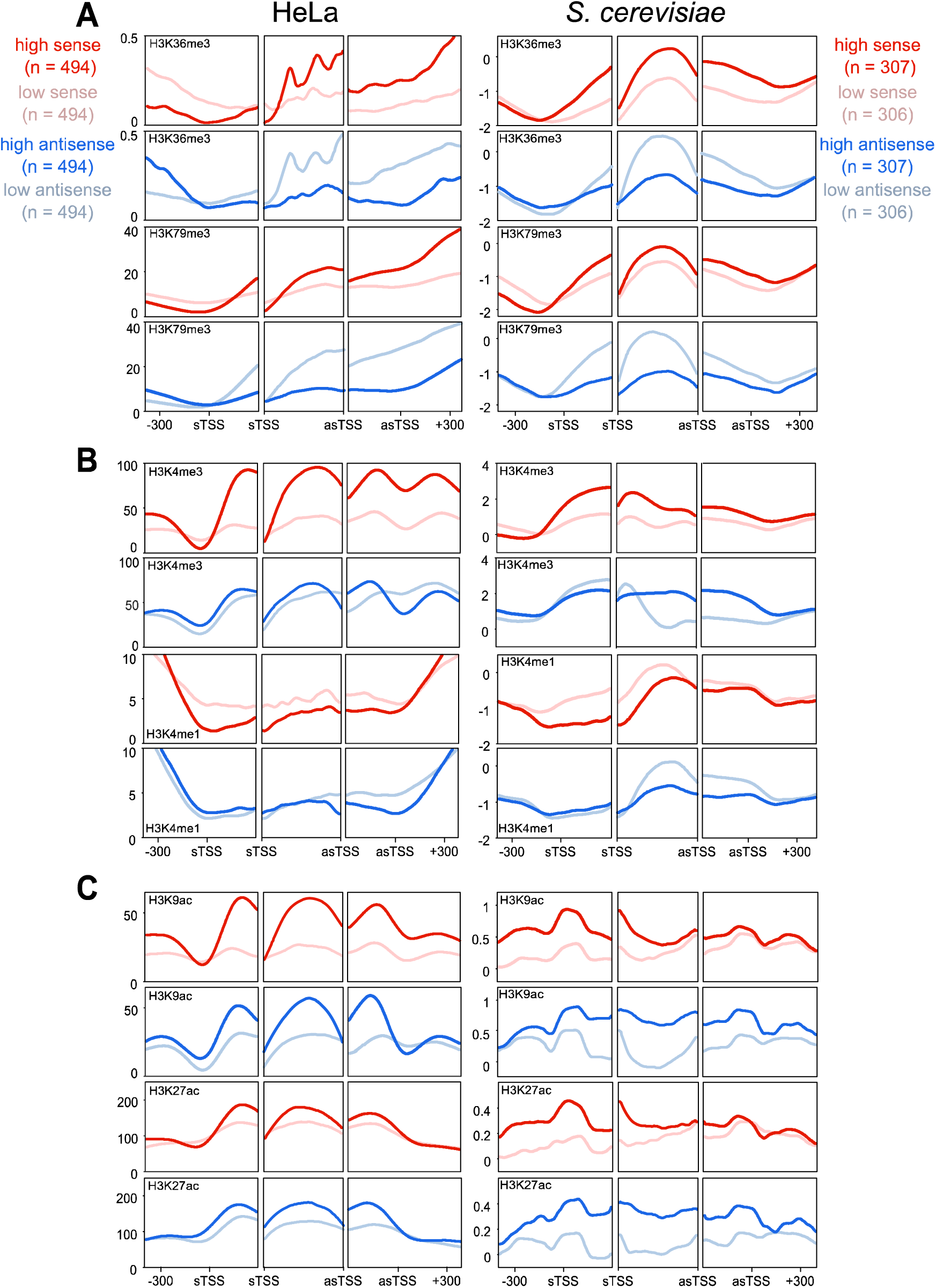
Antisense transcription has similar associations with chromatin modifications in both yeast and humans. (A) The average levels of H3K36me3 and H3K79me3 and antisense transcription in HeLa and *S. cerevisiae* genes. For each trio of panels, the left panel shows average levels around the sTSS, the right panel shows the average levels around the asTSS, and the middle panel shows the average level within thirty equal sized bins within the region bound by the sTSS and asTSS. Genes considered are selected from those that contained an asTSS, as defined in Fig 1. Shown in red are two sets of genes – those with high levels of sense transcription (dark red), and those with low levels (pale red), determined by GRO-seq in HeLa cells, and PRO-seq in budding yeast. Shown in blue are those genes with high levels of *antisense* transcription (dark blue), and those with low levels (pale blue), determined by GRO-seq in HeLa cells, and PRO-seq in budding yeast. **(B)** Average levels of H3K4me3 and H3K4me1, laid out as in **A. (C)** Average levels of H3K9ac and H3K27ac, laid out as in **A**. Relates to Figure 3.

**Supplemental Figure 3.**
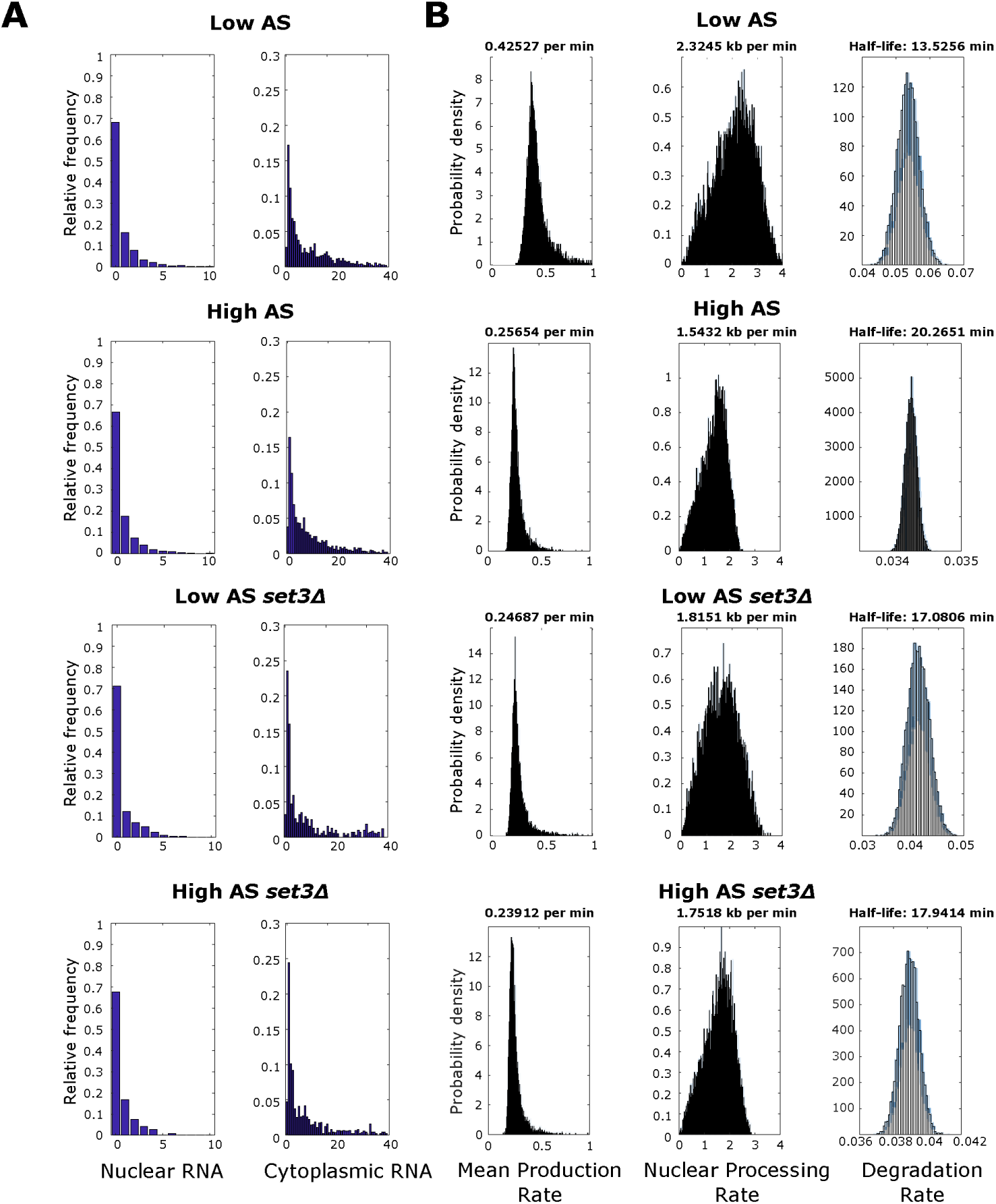
Nuclear and cytoplasmic RNA-FISH distributions for *GAL1* foci and dynamics of transcription and transcript processing. **(A)** The distribution of foci in the nucleus or cytoplasm from WT (top 2 panels) or *set*3Δ (bottom 2 panels) with High or Low antisense transcription, as indicated. **(B)** Plots showing the probability density for mean production rate (left panel), nuclear processing rate (middle panel) and degradation rate (right panel) for WT or *set*3Δ strains with High or Low antisense transcription, as indicated. The most likely rate is indicated above each plot. Relates to Figures 4-6.

**Supplemental Figure 4.**
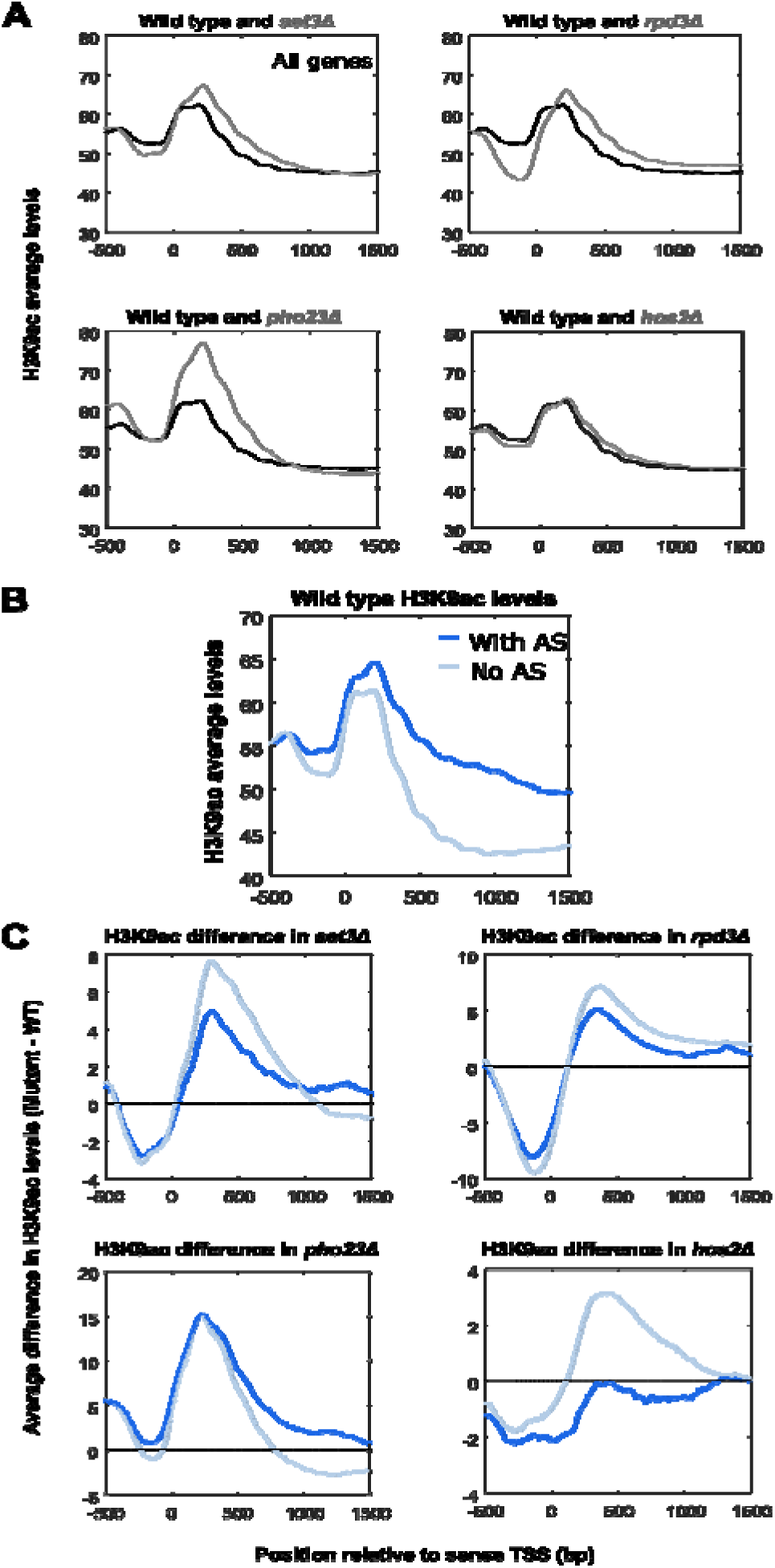
The presence of an antisense transcript can change how H3K9 acetylation levels changes following deletion of a histone modifying enzyme. **(A)** Average levels of H3K9 acetylation in all budding yeast genes, in wild type and mutant strains. **(B)** Average levels of H3K9 acetylation in wild type in two groups of yeast genes: those with an asTSS and those without, as defined in main text. **(C)** Average difference in H3K9 acetylation between mutant strains and wild type, for both genes with and without an asTSS. Relates to Figure 6.

**Supplemental Figure 5.**
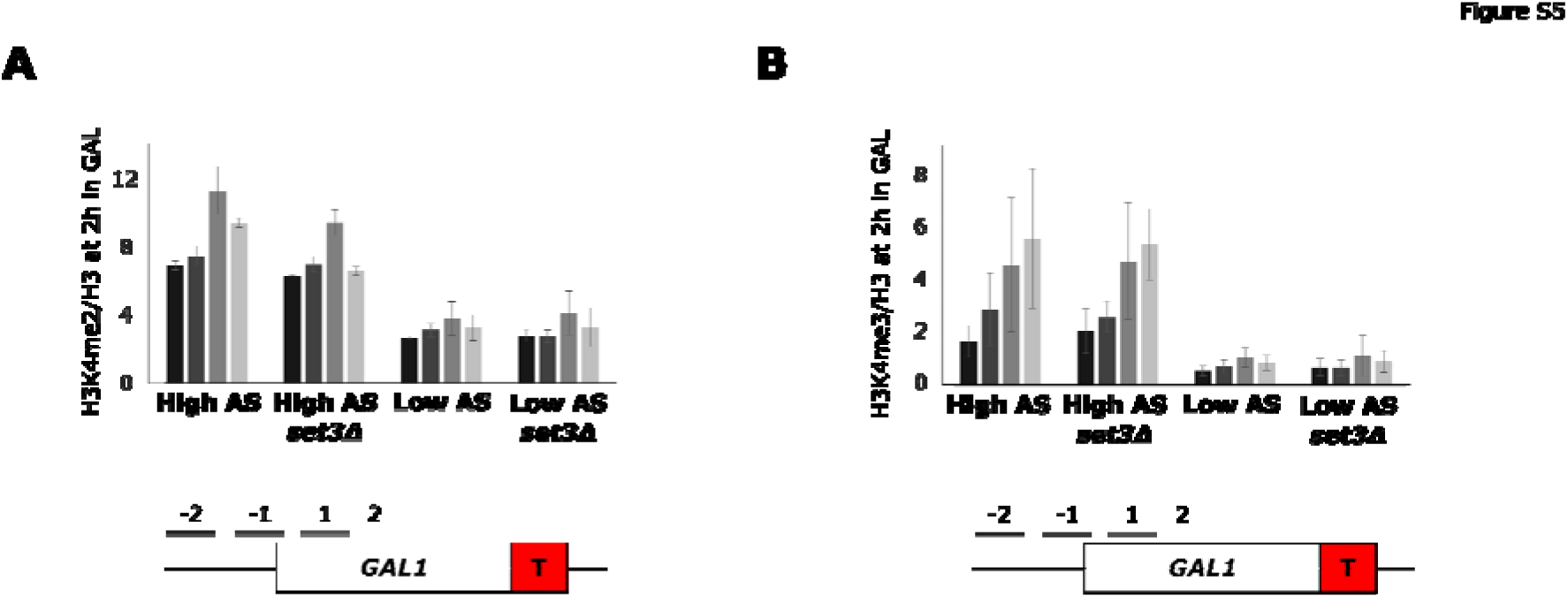
H3K4me2/3 does not change upon *SET3* deletion. **(A)** Levels of H3K4me2 relative to histone H3 at the engineered *GAL1* gene containing the altered ADH1 terminator (T) as measured by ChIP-qPCR at the primer positions indicated in the schematic below in the strains with high and low antisense in the presence and absence of *SET3*. N=2, error bars are SEM. **(B)** As in **A** but for H3K4me3. Relates to Figure 6.

## STAR Methods

## CONTACT FOR REAGENT AND RESOURCE SHARING

As Lead Contact, Jane Mellor is responsible for all reagent and resource requests. Please contact Jane Mellor at with requests and inquiries.

## EXPERIMENTAL MODEL AND SUBJECT DETAILS

All *Saccharomyces cerevisiae* strains used in this study are listed in the Key Resources Table. All strains and genetic manipulations were verified by sequencing or PCR-based methods.

## METHOD DETAIL

### Genetic manipulation of yeast strains

All strains used in this study are listed in the Key Resources Table. Genetic manipulation of strains was performed using the homologous recombination method ((Longtine et al., 1998). For gene deletion strains, PCR products were made containing the HISMX or KANMX selection cassettes flanked at both ends by 40 bp of sequence homologous to sequences either side of the region to be deleted. Construction of the *GAL1:ADH1*t (high AS) and TATA mutant (low AS) strains have been described previously (Murray et al., 2015; Murray et al., 2012). Cells to be transformed were grown to log phase, pelleted, re-suspended in 450 μl 100 mM LiAc/TE and incubated (> 1 h, 4°C). 100 μl of cell suspension, 10 μl of PCR product, 10 μl calf thymus DNA (Sigma D8661) and 700 μl 40% polyethylene glycol in 100 mM LiAc/TE were incubated (30 min, 30°C) then heat-shocked (20 min, 42°C). Cells were pelleted (5 min, 7,000 rpm), re-suspended in H_2_0 and plated onto appropriate selection media. DNA was extracted from the resulting colonies, screened by PCR and confirmed by sequencing.

### Yeast culture

Strains were streaked from glycerol stocks onto 2% agar YPD (1% yeast extract (Difco), 1% bactopeptone, 2% glucose) plates and grown (1-2 days, 30°C). Cell pre-cultures were then grown overnight in 5 ml YPD at 30°C. This culture was used to inoculate an appropriate volume of YPD culture at OD_600_ 0.2 which was grown at 30°C, shaking at 200 rpm to OD_600_ 0.45. To induce the *GAL1* gene, cell cultures were centrifuged (3,000 rpm, 3 min) and then re-suspended in YPG (1% yeast extract (Difco), 1% bactopeptone, 2% galactose) pre-warmed to 30°C. Re-suspended cells were incubated (30°C, 200 rpm) for the specified time(s) before harvesting by centrifugation (3,000 rpm, 4 min). For the experiments to obtain the *GAL1* sense degradation rates, after 2 h in YPG, cells were transferred back to fresh YPD pre-warmed to 30°C and 15 ml samples were harvested at 0, 5, 10, 20, 30 and 60 min.

### Chromatin Immunoprecipitation (ChIP)

Yeast grown to OD_600_ 0.5 in 50 ml of YPD were transferred to YPG for 2 h before they were fixed in 1 % formaldehyde in 45 ml PBS for 30 min at 22°C followed by addition of 125 mM glycine for 5 min. Cell pellets were collected by centrifugation (3,000 rpm, 4 min) before washing twice with 10 ml cold PBS. Cells were re-suspended in 500 μl cold FA-150 buffer (10 mM HEPES pH 7.9, 150 mM NaCl, 0.1 % SDS, 0.1 % sodium deoxycholate, 1 % Triton X-100) and broken using 1 ml glass beads on a MagnaLyser (Roche) (2 x 1 min runs, 6,000 rpm, 4°C). Sample volume was increased to 2 ml with FA-150 buffer before shearing of the fixed chromatin by sonication using a biorupter (Diagenode, 30 min, 1 min on, 20s off, medium setting). Chromatin was cleared by centrifugation (10,000 rpm, 15 min, 4°C) and 50 μl was diluted to 200 μl with FA-150 buffer and incubated with 5 μl of the following antibodies as appropriate: H3, H3K4me2, H3K4me3, H3K9ac (for details see the Key Resources Table) in 1.5 ml siliconised Eppendorf tubes for 15-20 h rotating at 4°C. Bound chromatin was immunoprecipitated for 90 min at 22°C with 50 μl protein A-Sepharose pre-blocked with bovine serum albumin and sonicated salmon sperm DNA. Beads and attached chromatin were pelleted by centrifugation (2,600 rpm, 1 min) and washed with TSE-150 buffer (20 mM Tris-Cl pH 8.0, 150 mM NaCl, 2 mM EDTA, 0.1 % SDS, 1 % Triton X-100) for 3 min, TSE-500 buffer (20 mM Tris-Cl pH 8.0, 500 mM NaCl, 2 mM EDTA, 0.1 % SDS, 1 % Triton X-100) for 3 min, LiCl buffer (0.25 M LiCl, 10 mM Tris-Cl pH 8.0, 1 mM EDTA, 1 % dioxycholate, 1 % NP-40) for 15 min and twice with TE. After washing, chromatin was eluted from the beads for 30 min at 65°C with elution buffer (0.1 M NaHCO3, 1 % SDS). Addition of 350 mM NaCl and incubation for 3 h at 65°C reversed the cross-links before treatment of samples with RNase A for 1 h at 37°C and proteinase K overnight at 65°C. DNA was purified using a PCR-purification kit (Qiagen) and eluted in 400 μl 1 mM Tris-Cl pH 8.0. Input DNA was diluted accordingly. Real-time quantitative PCR (qPCR) was performed using a Corbett Rotorgene and Sybr green mix (Bioline). Data ([IP - no antibody control]/input) were expressed as a percentage of the input and normalised to levels of H3 where appropriate. The primers used are listed in Table S1. All ChIP experiments were performed ≥2 times with independent biological samples.

### RNA extraction

15 ml of log phase yeast culture at a density of OD_600_ 0.6-0.8 grown in the appropriate medium was pelleted (3,000 rpm, 3 min), re-suspended in 400 μl TES (100 mM Tris-HCl (pH 7.5), 100 mM EDTA (pH 8.0), 0.5% SDS) and 400 μl phenol:chloroform (pH 4.7) and incubated (65°C, 20 min, 1,400 rpm). The mixture was incubated (-80°C, 30 min). After spinning (13,000 rpm, 20 min, 4°C), the upper layer was transferred to 10 mM NaOAc pH 5.5/ethanol and incubated (-80°C, >30 min). RNA precipitate was pelleted (13,000 rpm, 20 min, 4°C) and re-suspended in 100 μl H_2_0. RNA concentration was measured using a Nanodrop and samples were diluted to 1,000 ng/μl.

### Northern blotting

20 μg of RNA was separated on 1.1% formaldehyde FA gels for 3 h and transferred to Hybond-N+ nylon membranes (Amersham) by wet blotting overnight in 20xSSC. After fixing the RNA to the membrane (2 h, 80^o^C), the membranes were blocked in PerfectHyb Plus (Sigma H7033) (>2 h, 65^o^C). Radio-labelled strand-specific *GAL1* sense and *GAL1* antisense probes were generated using asymmetric PCR with the primers listed in Table S2. After probe purification with in-house-constructed Sephadex G-50 columns, the probe was added to the tubes containing the membranes and hybridised overnight at 65^o^C. Non-specifically bound probe was removed by washing the membranes twice in 1xSSC/0.1 % SDS and once in 0.2xSSC/0.1 % SDS, 0.1xSSC/0.1 % SDS and 0.05xSSC/0.1 % SDS for 20 min each at 65^o^C. Membranes were typically exposed to X-ray film for 1 h-1 week. For quantification, images were acquired using a FLA 7000 phosphorimager (GE Healthcare). Levels of the 18S and 25S rRNA species measured by ethidium bromide staining were used as loading controls. All Northern blotting experiments were repeated ≥2 times with independent biological samples.

### RNA fluorescence in situ hybridization (RNA-FISH)

50 ml yeast culture was grown in YPD to >0.45 OD_600_ before transfer to YPG for 2 h. 50 ml cells at OD_600_ 0.6 were pelleted (3,000 rpm, 4 min) and fixed with 4% (v/v) paraformaldehyde in PBS (45 min, 80 rpm, 22°C). Fixed cells were washed twice with 10 ml FISH buffer A (1.2 M sorbitol, 0.1 M KHPO_4_ (pH 7.5)) and re-suspended in 1 ml FISH buffer B (FISH buffer A, 20 mM ribonucleoside vanadyl-complex (VRC), 20 μM 2-mercaptoethanol). The mixture was incubated (15-40 min) at 30°C with 15 μl lyticase (25 U μl^−1^, Sigma) until > 70% of cells were spheroplasted, as observed by microscopy. Cells were pelleted (3,000 rpm, 3 min, 4°C) and washed with and then re-suspended in 1 ml FISH buffer B without 2-mercaptoethanol. ~150 μl of cells were left to settle (30 min, 4°C) on poly-L-lysine treated coverslips. These were gently washed with 2 ml FISH buffer A to remove unattached cells and incubated (-20°C, > 3 h) in 2 ml 70% ethanol. Samples were rehydrated twice with 2 ml of 2X SSC for 5 min at room temperature and washed with 40 % formamide in 2xSSC. For the hybridization, 0.5 ng of each probe, 10 μg *E. coli* tRNA and 10 μg salmon sperm DNA were mixed and lyophilized in a SpeedVac. 12 μl of 40% formamide, 2X SSC, NaHPO_4_ pH 7.5 was added and the probes were denatured at 95°C for 3 minutes followed by the addition of 12 μl of 2xSSC, 2 mg/ml BSA, 10 mM VRC. Hybridization was performed overnight at 37°C in a parafilm-sealed chamber, where the coverslips with the cells facing down were placed onto 22 μl of the hybridization mixture. The coverslips were then subjected to a series of washes: twice with 40% formamide/2xSSC (15 min, 37°C); once with 2xSSC, 0.1% Triton X-100 (15 min, 22°C); once with 1xSSC (15 min, 22°C); and once with 0.05xSSC (15 min, 22°C). The coverslips were dipped into H_2_O. Once dry, coverslips were mounted onto a microscope slide using ProLong Diamond Antifade Mountant with DAPI (Life Technologies), allowed to polymerize for 24 h in the dark and then sealed with nail varnish. Cells were imaged using a DeltaVision CORE wide-field fluorescence deconvolution microscope using a 100x/1.40 objective lens. 21-31 0.2 μm z stacks were imaged with an exposure time of 0.01 s and 1 s for DAPI and Cy3 channels respectively. All RNA-FISH experiments were repeated ≥2 times with independent biological samples.

### RNA-FISH probe design and synthesis

DNA probes of ~50 nt and ~50 % GC content were designed with five modified bases (amino-allyl dT) spaced by about ~10 nt included for the incorporation of the fluorophore (see Table S3 for probe sequences). Modified DNA oligos were custom ordered from MWG Eurofins. For the labelling of the probes, a total of 5 μg was purified using the QIAquick Nucleotide Removal Kit (Qiagen) and eluted with 40 μl of H_2_O. The probes were then lyophilized in a SpeedVac, resuspended in 10 μl of 0.1 M sodium bicarbonate pH 9.0 and added to the dye-containing tube (CyDye™ GE Healthcare, Cy3 PA23001). The tube was vortexed vigorously followed by a quick spin. The reaction was incubated overnight at room temperature with low speed shaking. The probes were purified using the QIAquick Nucleotide Removal Kit (Qiagen) and eluted with 100 μl of elution buffer (supplied with the kit). The concentration and efficiency of the labelling was measured using a spectrophotometer. Probes were stored in the dark at −20°C. The labelling efficiency was calculated as described (Zenklusen and Singer, 2010).

## QUANTIFICATION AND STATISTICAL ANALYSIS

### RNA FISH analysis

Image quantification was performed using custom Matlab (MATLAB Statistics and Image Toolboxes Release 2015a, The MathWorks, Inc., Natick, Massachussetts, United States) scripts based in part on elements of FISH-quant (Mueller et al., 2013) and CellProfiler (Carpenter et al., 2006), and utilising MIJI (https://imagej.net/Miji) and MIJ (http://bigwww.epfl.ch/sage/soft/mij/) to import data from FIJI (Schindelin et al., 2012). The custom scripts allowed for greater automisation of the quantification process than is possible with FISH-quant and the algorithms were tailored to our data.

#### Deconvolution and background subtraction

Images were deconvolved with a conservative deconvolution method and 10 cycles using DeltaVision Softworx software. DAPI and Cy3 channels for the images were processed separately. Images were background-corrected with the following procedure. The median of all pixel intensities for each channel and each image, *p*_*med*_, was found. This was chosen as it was observed to generally be close to the modal value of the distribution. A measure of the spread around this value was also found by constructing a metric similar to the standard deviation from all pixels with intensities less than or equal to the median intensity. The median plus this spread value was taken to be the background value and subtracted from all pixels, background = *p*_*med*_ + sqrt[(1/(*N*_i_-1)) sum_*i*_((*p*_*i*_ – *p*_*med*_)^2)], where *i* runs over pixels with intensity less than or equal to *p*_*med*_, *N*_*i*_ is the number of pixels with intensity less than or equal to *p*_*med*_ and *p*_*i*_ is the intensity of pixel *i*. Thus, the new intensity of *p*_*j*_ was *p*_*j*_ – background where *j* runs over all pixels. Any pixels that had negative intensity following this were set to 0.

#### Foci identification and thresholding

Candidate foci in the FISH (Cy3) channel were initially identified using Piotr’s Matlab toolbox (https://pdollar.github.io/toolbox) nonMaxSupr function with a 1 pixel radius for detection. Images from each biological repeat and strain were processed together. To distinguish foci from random clustering of fluorescently-labelled molecules, foci intensities were compared between the strains under testing and a double knockout strain, *gal10-1*ΔΔ, which has no sequences to which the FISH probes should hybridise. In each experiment, histograms of all foci intensities as returned by the nonMaxSupr algorithm were constructed (bin width 250, normalised by probability). For each strain a tentative intensity cut off was taken to be the first bin in which there was 10 times more signal in the strain than the knockout (with manual adjustment for obvious outliers). Within each experiment (which could contain multiple strains) the final cut off value was taken as the mean of the tentative cut-off values. All foci with intensities less than the cut-off value from a set were not considered in all further analysis.

The deconvolution, background subtraction and foci identification are performed by the supplied FindAndAnalyseFoci Matlab function. This function should be run on all strains and the knockout from a single experiment before a threshold for valid foci is determined. This threshold can then be found using the DetermineCutoffs Matlab script.

#### Automated nuclei detection

A separate script automatically identifies nuclei and cells and quantifies the foci that fall within the nuclear and cellular boundaries. The first step of the process is to identify the nuclei in 3 dimensions using the DAPI channel. The procedures followed here allowed for improved detection of individual nuclei that differed in brightness or were very close to neighbouring nuclei. The DAPI channel of each image was scaled to the minimum and maximum intensity pixels, i.e. *p*_*i*_ = ( *p*_*i*_ – min(*p*)) / ( max(*p*) – min(*p*)), where *i* runs over all pixels, min(*p*) and max(*p*) denote the minimum and maximum pixel intensities of the set respectively. The scaled DAPI images then have a Gaussian filter applied using Matlab function imgaussfilt3 with a smoothing-kernel standard-deviation value of 2. For each processed image Otsu’s method for multiple thresholds, Matlab function multithresh, was used to give 6 threshold levels and the image was segmented into seven levels around these using the Matlab function imquantize. The segmented images thus contained pixels with values from 1 to 7. Each segmented image was then restricted to a subsection of the available z-stacks by setting any pixels in z planes below or above certain values to zero, to avoid any errors due to using overly blurred portions of the image. For images with 31 z stacks images were typically restricted to include only pixels from z planes 12 to 22, inclusive, and for images with 21 z stacks images were typically restricted to z planes from 2 to 20, inclusive. These values were manually adjusted in some cases to allow for off centred focusing but the same planes were used for all images of a particular strain taken in a single experiment.

The segmented levels were cycled through from the 3^rd^ to the 7^th^ levels. For each level, a 3D logical image was formed from the pixels with value equal to the value of the level. The z stacks were then cycled through and all holes (areas with pixels with value 0 inside areas with value 1) were filled, Matlab function imfill with flag ‘holes’. Then, all 3D-contiguous regions (with 26 connectivity) of pixel value 1 that had more than 6000 or fewer than 50 pixels were removed by setting all pixels in the region to zero, using Matlab xor and bwareaopen functions. Any remaining 3D-contiguous regions (26 connectivity) were then labelled using the Matlab function bwlabeln. For levels lower than the final level, an identical procedure was performed on the level immediately above, without the final labelling step. Each labelled section was then cycled through and if there was no overlap with any non-zero valued pixels in the segmented level above, it was deemed as a good candidate for a nucleus and saved (the centroid was determined with the Matlab function regionprops and flag ‘centroid’ and each coordinate was rounded to the nearest integer). Any overlap with the level above signified the existence of a superior candidate or superior candidates in this region and this potential nucleus was not saved. This process was repeated until the final level in which no checking against a higher level was possible.

Once all good nuclear candidates had been identified in this way, any nuclei that were very close together were merged with the following procedure. The Euclidean distances, measured in pixel coordinates, between all centroid locations were calculated. A list of all non-equal pairs of centroids that had a distance of less than or equal to 6 between them was created. This was done by having two nested loops: the outer loop cycled over the centroids from *i* = 1 to (*N*_*c*_-1) and the inner loop cycled over *j* = (*i*+1) to *N*_*c*_. Any *i, j* pairs from this loop with distance less than or equal to 6 were listed. If any centroid appeared more than once in the list, this list was reordered in ascending order of distances. The ordered list was then cycled through in order and, starting with the *i* element of the pair, if this centroid was repeated, all but this first appearance of this centroid in the list were deleted and then the same test and deletion was done with the second centroid of the pair. Then all pairs of centroids on the list were merged by taking the average of their coordinates and rounding each coordinate to the nearest integer. The distances between the new centroids were calculated and the process was repeated until no centroids that were within a distance of 6 from another remained. This procedure prioritised merging centroids that were closest together in the case that there were multiple possible mergers.

The list of centroids generated in this way was then used to generate 3D masks of the nuclei with the following procedure. The centroids were cycled through and the mean intensity of the 27 pixels surrounding and including the centre pixel was taken, using the filtered and z-stack restricted DAPI signal. Logical images were formed for each centroid by setting all pixels with intensity greater than or equal to 0.65 times this mean value to unity and all others to zero. The 26-connected 3D component from this that overlapped with the centroid position was taken as the 3D nuclear mask for this centre point. To avoid counting areas that were too low intensity relative to the image, nuclei that had a mean intensity of the 27 pixels less than 0.025 were rejected. Once this list of nuclear masks had been created, a further filtering was done by removing any nuclei that had a volume of fewer than 50 pixels. For later analysis, additional 2D nuclei masks were formed by taking the maximum of the 3D nuclei through the z stacks which due to the masks being stored as logical images corresponds to the greatest extent in the x and y coordinates that the nucleus has in any of the allowed z stacks. At this step the 2D nuclei were relabelled based on connected components with 8 connectivity (Matlab function bwlabel) as it is possible for 3D nuclei to not touch but to overlap when flattened in this way. For later classification of foci, these 2D nuclear centres were extruded to fill the allowed z stacks.

#### Automated cell detection

The 2D nuclei identified above were used as seed points to identify cell outlines. Cell masks were identified using the CellProfiler function IdentifySecPropagateSubFunction from the MEX compiled file supplied with the developer’s version of CellProfiler 1.0. Cells were identified with a combination of the DAPI signal, FISH foci signal and autofluorescence of the cells observed in the FISH channel. The identifySecPropagateSubFunction takes a number of inputs: a set of seed points for cells; a 2D image with varying intensity; a 2D logical image; and a regularization factor which determines how to weight between the 2D images and distance to the nearest seed points when determining cell boundaries. A regularization factor value of 0.0001 was used in all cases.

The 2D images with varying intensities were constructed by combining processed DAPI and FISH channels in the following way. For each channel of each image, the sum in the z direction was taken to get a flattened image and the maximum and minimum intensities observed in this image were found. Each pixel in the image was then normalised as *p*_*i*_ = ( *p*_*i*_– min(*p*)) / ( max(*p*) – min(*p*)). For each image the normalised flattened channels were averaged and then this average had a Gaussian filter applied (Matlab function imgaussfilt with smoothing-kernel standard-deviation value of 2).

The 2D logical images were constructed starting in a similar way by creating normalised and flattened images as above prior to the averaging step. Each channel for each image had a Gaussian filter applied but the DAPI signal used a smoothing-kernel standard deviation of 5 and the FISH channel used a value of 2. Each filtered channel for each imaged then had a threshold generated by taking the lowest value from a two-level Otsu’s-method thresholding (Matlab function multithresh with 2 levels). Each of the filtered channels for each image was then converted into a logical image with pixels having an intensity less than the threshold being set to 0 and the rest being set to 1. The channels for each image were then combined using the logical OR operation.

The nuclei and cells were then further processed to remove anything touching a border of the image as cells touching the border are likely to have part of the cell out of the image and using them for data collection could bias the results. First any 3D-resolved nuclei that touched the border in 3D were removed (Matlab function imclearborder). Following this, any cell masks that were touching the border were removed. Then any 3D nuclei considered as being too large (i.e. containing more than 3000 pixels) were removed. The size-based removal was done after cell detection as large detected nuclei often corresponded to multiple nuclei close together and keeping these causes the corresponding cells to be detected as one large cell, which often aided in assigning the correct boundaries to nearby cells. Then any cells that did not overlap with any nuclei were cleared and any nuclei that did not overlap with a cell were also cleared. Finally, any cells that had 2 or more nuclei were removed along with the corresponding nuclei. This case can happen rarely when 3D nuclei do not touch but overlap when collapsed onto 2D resulting in a merged nucleus. These were removed as it is likely that there would be some cytoplasmic overlap also in this case.

#### Foci classification and mRNA quantification

Cells were first divided into nuclear and cytoplasmic components. The nuclear components were formed first by flattening the 3D-resolved nuclei masks (taking the maximum over the z-stacks) and then extruding them to the allowed z stacks (the same z stack limits used when restricting in the nuclei-detection phase). The 2D cell masks were then extruded to the same allowed z stacks and had the nuclear area within them set to zero, which gave the cytoplasmic components. Foci whose centre pixel lay in the nuclear region where classified as nuclear transcripts and foci whose centre pixel lay in the cytoplasmic region were classified as cytoplasmic transcripts. Any foci lying outside these regions were discounted from all further analysis.

In order to quantify the foci in terms of RNA molecules an intensity value for all accepted foci was calculated by taking the mean of the 27 pixels immediately surrounding and including the central pixel (found by rounding each coordinate of the output of the nonMaxSupr function to the nearest integer). For a strain and image set from a single experiment, the median of these intensity values was calculated and was taken to correspond to a single RNA molecule. There is no reason that a FISH focus should contain only a single RNA, especially in the nucleus, but we assumed that foci most commonly contained a single RNA. Each focus intensity was then converted into a corresponding number of mRNA molecules by dividing by the median intensity and rounding to the nearest integer. Note that this can result in foci being classified as containing zero RNA molecules and provides an additional filtering step similar to the initial cut off based on knockout strains.

The detection of nuclei and cells, and the quantification of foci is performed by the supplied Matlab script DetectCellsAndQuantifyFoci. This script should be run after the FindAndAnalyseFoci function and DetermineCutoffs script as it uses the data generated in the first script and the cut off generated after averaging the determined cut offs over an experiment. The cut off must be manually changed in the DetectCellsAndQuantifiFoci script before running. Scripts are available from doi:10.17632/dhnvj4xs5d.1.

### Mathematical Modelling

RNA synthesis and degradation were modelled as described in the main text. 4 of the 5 parameters were sampled via Latin Hypercube with 1,000,000 sampling points. The degradation rate was sampled as described below. 10,000 cells were simulated for 500 min to reach steady state and the number of nuclear and cytoplasmic RNA recorded. The Kolmogorov-Smirnov statistic was used as a goodness-of-fit metric to compare simulated results to raw data. The best 10,000 parameter sets as judged by fit to nuclear and cytoplasmic RNA were then taken forward. For each strain, the modal value of the histogram of the mean initiation rate (on*init/(on+off)) was taken from the fits to the cytoplasmic data. The values for the nuclear processing rate were then determined by sampling data points from the ratio of mean initiation rate to nuclear export rate giving the fits to the nuclear data. Again, 10,000 parameters were sampled from the values determining the nuclear distributions, following the mean initiation rate distribution given by the cytoplasmic data.

### Chromatin Immunoprecipitation

Real-time quantitative PCR (qPCR) was performed using a Corbett Rotorgene and SYBR green mix (Bioline). qPCR was performed in triplicate for each sample and quantified using a standard curve. Histone modification data ([IP - no antibody control]/input) were expressed as a percentage of the input and normalised to levels of histone H3 at each amplified region. Data are presented as averages of ≥2 biologically independent experiments, with error bars representing the standard error of the mean.

### Northern blotting

Raw images for quantification were captured using a FLA 7000 phosphorimager (GE Healthcare). Mean intensity of band was quantified using Fiji/ImageJ. Normalised levels of RNA were obtained from Northern blots and a degradation rate was obtained by fitting results to an exponential using MATLAB fit function. Root mean square error was taken as goodness-of-fit metric and correspondingly used as the standard deviation of the estimator of the exponential decay term. To sample degradation rates for the purposes of modelling, degradation rates were sampled from a Beta distribution with maximum and minimum values given by the 95% confidence intervals of the estimator with standard deviation equal to the root mean square error of the estimator.

### Bioinformatic analysis

#### Identification of sense and antisense TSSs in yeast and humans

Cap analysis of gene expression (CAGE) data in HeLa cells was obtained from the ENCODE repository on the UCSC Genome Browser (Rosenbloom et al., 2010), and used to determine genome coordinates of sense and antisense transcript start sites (sTSSs and asTSSs respectively). Data were pooled from nuclear and cytoplasmic fractions, both polyadenylated and non-polyadenylated, and from whole cell extract, for which only polyadenylated data were available. CAGE cluster coordinates, determined with an HMM algorithm applied to the CAGE tag data, were obtained from the same source. To determine TSS coordinates in HeLa, we took the same approach presented previously by (Conley and Jordan, 2012). Clusters were ignored if they contained less than 2 overlapping CAGE tags, as it has been previously reported that two or more overlapping tags represent validated TSSs (Carninci et al., 2006; Faulkner et al., 2009). The TSS coordinate of a given cluster was taken to be the base with the highest density of mapped CAGE 5’-ends. As an added step, TSSs were excluded from all subsequent analyses if they contained less than 3 NET-seq reads in a 200 bp window immediately downstream, within the same orientation. To determine the sTSS of a given protein-coding gene, we scanned within a region 500 bp upstream of the left-most annotated TSS, and 500 bp downstream of the right-most annotated TSS. In the case of multiple sTSSs the one with the highest CAGE density was taken to be the predominant sTSS. asTSSs were determined by scanning between the sTSS and the annotated transcript end site, again, the asTSS with the highest CAGE density was considered the predominant asTSS in the case of multiple candidates.

Budding yeast sTSSs and asTSSs were determined using transcript isoform sequencing (TIF-seq; (Pelechano et al., 2013), using the list of major transcripts provided. For each gene, the sTSS was derived from the sense transcript which had the highest number of supporting reads in YPD, and which completely encompassed the open reading frame. The asTSS was taken as the antisense transcript with the highest number of supporting reads, and which overlapped the open reading frame in the antisense orientation.

To assess whether asTSSs were better aligned to the sTSS or the end of the 1^st^ exon in HeLa, we determine for each gene the distance between the sTSS to the asTSS, and expressed it as a fraction of the distance between the sTSS and the end of the 1^st^ exon. We compared the resultant histogram to a randomly generated distribution, in which the asTSS for each gene was randomly reassigned to a base within the region shown in Figure 1D. This approach was repeated in HeLa using the end of the 2^nd^ exon in place of the 1^st^, to assess whether asTSS showed preferential alignment to the 2^nd^ exon over the sTSS. It was also repeated in yeast, using the 3’ end of the open reading frame in place of the end of the 1^st^ exon (Figure 1F).

#### Correlating sense and antisense transcription

NET-seq data were obtained in HeLa cells from (Nojima et al., 2015), specifically their data obtained using an antibody against all forms of Pol II, phosphorylated and phosphorylated. NET-seq data in yeast were obtained from (Churchman and Weissman, 2011). To compare sense transcription levels between genes with and without an asTSS, we calculated the average number of NET-seq reads per base pair within the 1^st^ exon, and compared the distribution between the two gene groups using a Wilcoxon rank sum test. Correlations between the transcription levels of different sorts of transcript were calculated by determining the Spearman correlation coefficient between the numbers of NET-seq reads in the 300 bp windows shown in Fig 1B-D. The same approach was taken with the GRO-seq (for HeLa) and PRO-seq data (for yeast), obtained from (Core et al., 2014) and (Booth et al., 2016) respectively.

#### Assessing ChIP levels around sense and antisense TSSs

Genome-wide levels of histone-modifications and nucleosome occupancy were obtained from the following sources: For budding yeast, genome-wide levels of H3K36me3 and H3K79me3 were from (Kirmizis et al., 2009) (GSE14453). Levels of H3K4me1 and H3K4me3 were from (Kirmizis et al., 2007) (GSE8626). Levels of H3K9ac and H3K27ac were from (Weiner et al., 2015) (GSE61888). Nucleosome occupancy levels were from (Kaplan et al., 2009) (GSE13622). Genome-wide levels of gene compaction, determined using Micro-C, were from (Hsieh et al., 2015). Levels of H3K9ac in deletion strains of various histone modifying enzymes was from (Weinberger et al., 2012) (SRA051855.1). For HeLa cells, genome-wide levels of H3K36me3, H3K79me3, H3K4me1, H3K4me3, H3K9ac and H3K27ac were obtained from the ENCODE experiment matrix. Nucleosome occupancy levels were from (Kfir et al., 2015) (GSE65644). Levels of H3 histone modifications are not normalised to levels of H3. We assessed average levels only in genes with an asTSS, comparing the two quintiles with the highest and lowest levels of antisense transcription (determined by NET-seq) in a 300 bp window placed immediately downstream of the sTSS. We wished to simultaneously assess levels upstream of the sTSS, downstream of the asTSS, and in the region between both TSSs. To account for the varying distances between sTSS and asTSS, we broke this region into a hundred bins, calculating the average ChIP level within each bin.

#### Comparing gene compaction levels

Gene compaction levels in budding yeast were obtained from (Hsieh et al., 2015). Different gene groups were compared as discussed in the Results.

## DATA AND SOFTWARE AVAILABILITY

Unprocessed microscope images and analysis codes will be available on Mendeley

---

**Table S1.**
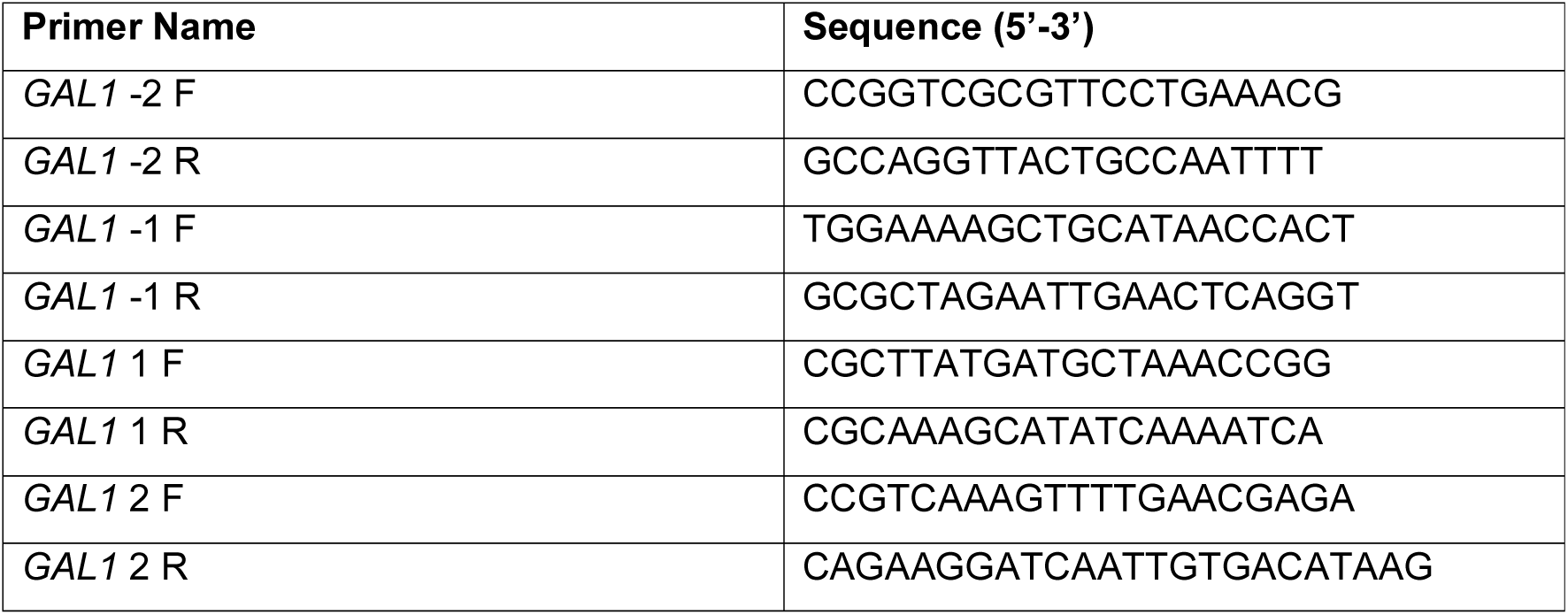
Primer sequences used for qPCR.

**Table S2.**
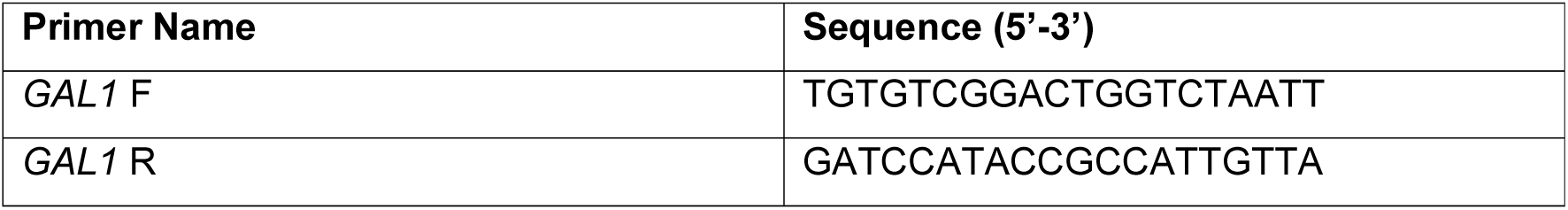
Primer sequences used to generate the strand-specific Northern blotting probes

**Table S3.**
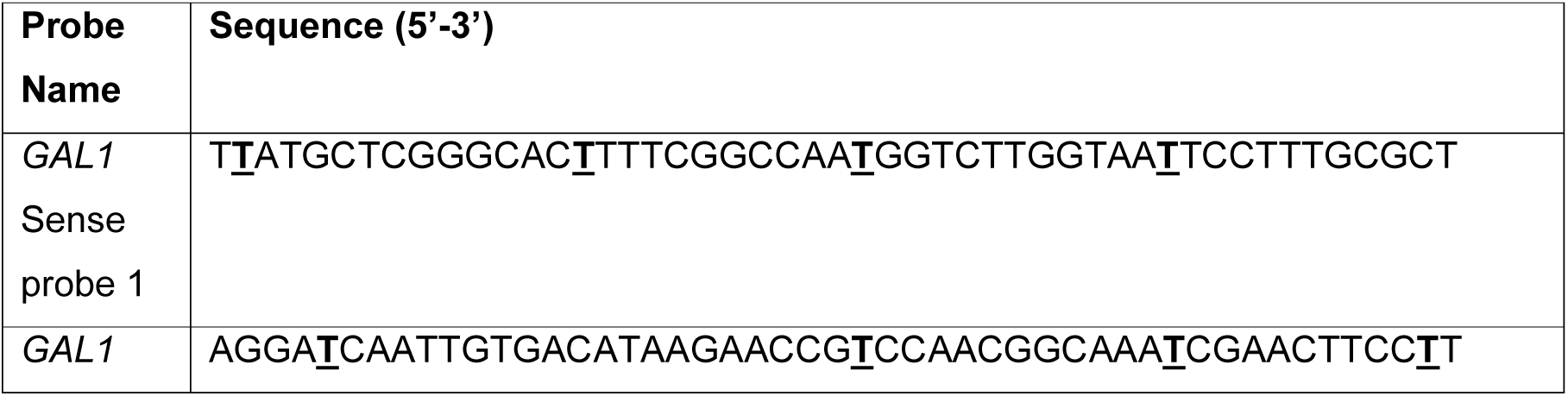
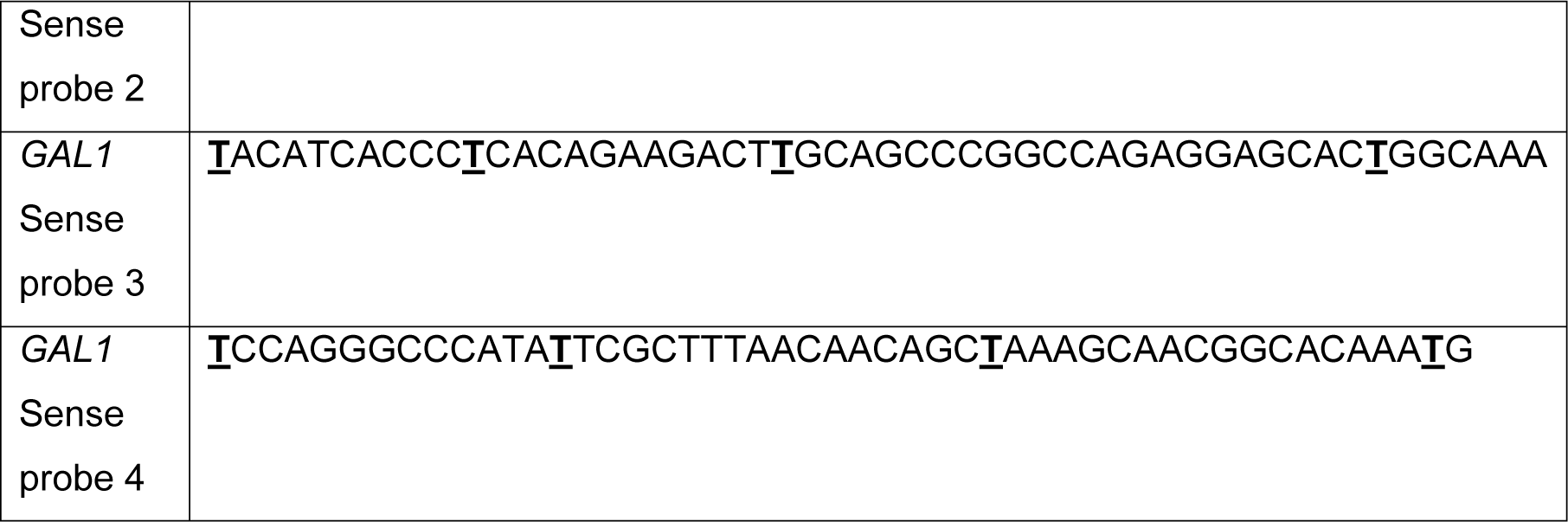
Sequences of the *GAL1* sense RNA-FISH probes. Underlined nucleotides were ordered as amino-allyl-dTs for subsequent labelling with Cy3 fluorescent dye.

**Table S4.**
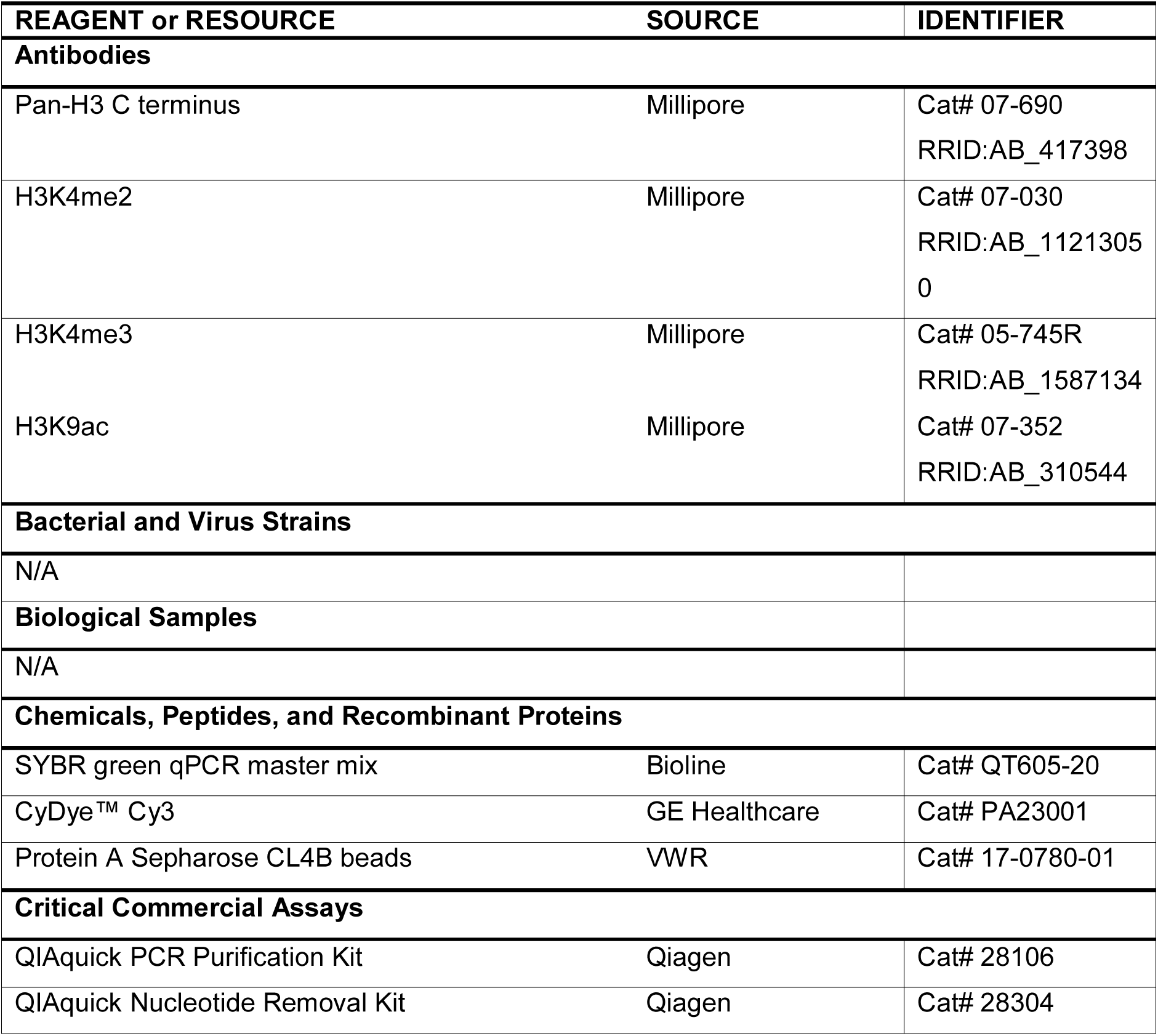
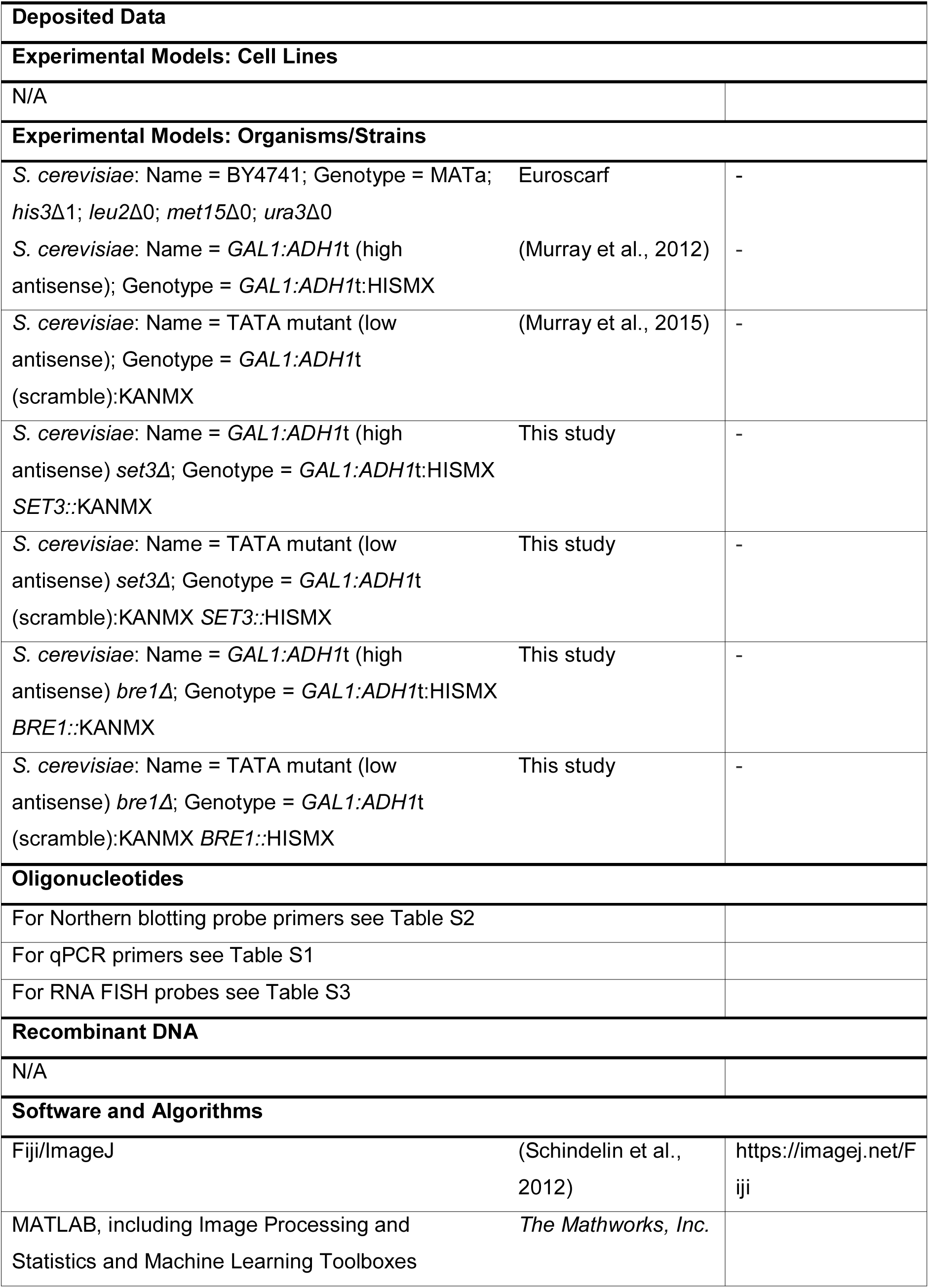
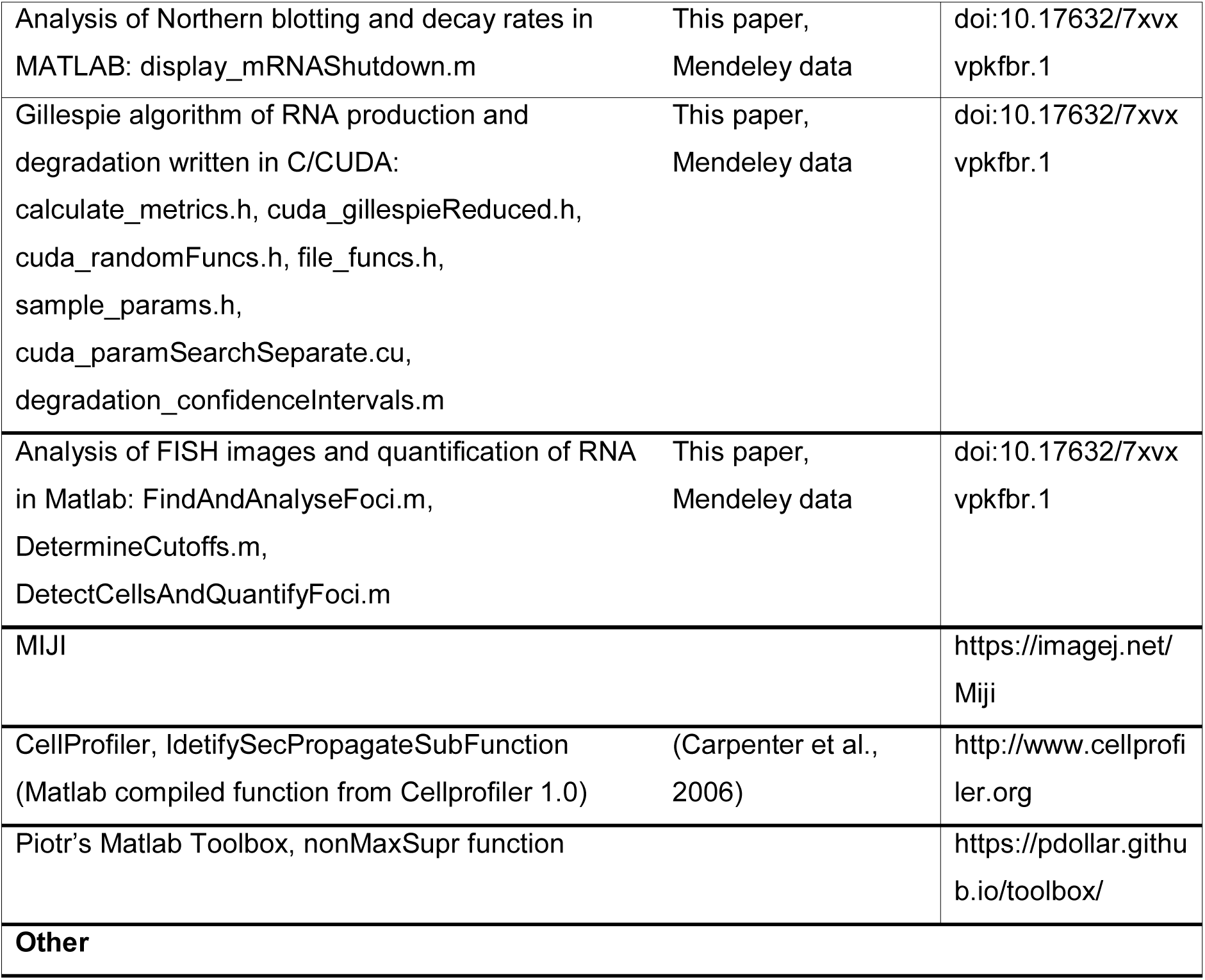
KEY RESOURCES TABLE

